# The *PERPETUAL FLOWERING* Locus: Necessary But Insufficient for Genomic Prediction of Runnerless and Other Asexual Reproduction Phenotypes in Strawberry

**DOI:** 10.1101/2025.04.20.646347

**Authors:** Hillel Brukental, Marta L. Bjornson, Dominique D.A. Pincot, Michael A. Hardigan, Sadikshya Sharma, Nicolas P. Jimenez, Randi A. Famula, Cindy M. Lopez Ramirez, Glenn S. Cole, Mitchell J. Feldmann, Steven J. Knapp

## Abstract

Strawberry (*Fragaria* × *ananassa*) reproduces sexually through seeds and asexually through stolons. The ability to cost-effectively clonally propagate hybrid individuals on a large scale has profoundly shaped strawberry breeding and production practices. Despite the technical and economic importance of clonal propagation, little is known about the genetic regulation of runnering in strawberry, other than the photoperiod-dependent pleiotropic effects of *PERPETUAL FLOWERING* (*PF*), a dominant, yield-doubling gene introgressed from a wild relative that knocks out temperature-dependent photoperiod sensitivity and partially suppresses runnering. Herewe showthat runnering phenotypes are highly variable and heritable, ranging from runnerless to prolific and unrestrained in both short-day (*pfpf*) and day-neutral (*PF*_) plants. We physically mapped the *PF* locus to Mb 26.4-27.3 on chromosome 4B using high-density SNP haplotypes and historical recombination among 932 individuals. *PF* was the only runneringassociated locus uncovered by genome-wide association studies among diverse clonal genetic resources and progeny from narrow and wide crosses (1,537 individuals). However, *PF* explained only 22% of the genetic variance for runnering. Genomic-estimated breeding values for runnering were accurately estimated especially among runnerless to near-runnerless individuals. Genomic prediction accuracies were greater within than between populations, increased when corrected for *PF*, and are sufficient for implementing genomic selection. These results pave the way for enhancing the productivity of strawberry by creating runnerless cultivars for seed-propagation and near-runnerless and reduced runnering cultivars for clone-propagation through phenotypic or genomic selection.

**CORE IDEAS:**

- Genetic variation for runnering is substantial and heritable
- The dominant *PERPETUAL FLOWERING* (*PF*) allele does not uniformly or ubiquitously suppress asexual reproduction
- Substantial wild donor DNA has persisted in day-neutral and summer-plant cultivars carrying the ’Wasatch’ *PF* allele
- The accuracy of *PF*-corrected breeding values for runnering are sufficient for implementing genomic selection
- Strawberry productivity can be improved by introducing cultivars with runnerless or near-runnerless phenotypes

## 2 INTRODUCTION

Cultivated strawberry (*F.* × *ananassa*), a self-compatible allo-octoploid (2n = 8x = 56), reproduces sexually through either self- or cross-pollination and asexually through stolons (runners) and plantlets (daughter plants) (Heide et al., 2013; Hytönen and Kurokura, 2020; Voth and Bringhurst, 1990). That reproductive flexibility has shaped breeding and cultivar development practices, which rely on the clonal propagation of parents and progeny (Darrow, 1966; Voth and Bringhurst, 1990). Strawberry cultivars are nearly always clonally propagated hybrid (F_1_) individuals originating from crosses between outbred parents with shared ancestry and complex, intertwined pedigrees (Darrow, 1966; Pincot et al., 2021). Shared ancestry, founder effects, finite populations, and strong directional selection have increased inbreeding in modern populations (Hardigan et al., 2021; Pincot et al., 2021; Feldmann et al., 2024b,a). Contrary to Porter et al. (2023), half- and full-sib matings have not been purposefully or strategically avoided in strawberry breeding. They are not only ubiquitous in strawberry pedigrees but commonly arise between parents with shared and often very close ancestry that inevitably and inexorably increase inbreeding (Robertson, 1961; Smith and Haigh, 1974; Wray and Thompson, 1990).

While breeding has traditionally and necessarily focused on improving fruit production traits in strawberry, adequate runner growth and clone yield (nursery production traits) are necessary for the cost-effective clonal propagation of cultivars, especially those grown on a large-scale for annual production systems. Uneconomical clone yields have generally not been a problem for modern short-day (SD) or day-neutral (DN) cultivars because most runner sufficiently in nursery production environments; however, some runner weakly, particularly modern DN cultivars purposefully selected for reduced runnering in fruit production environments, e.g., the reduced runnering day-neutral cultivar ’UCD Moxie’ (US Plant Patent #2019/0380247 P1).

Despite the economic and agricultural importance of asexual reproduction in strawberry, stolon and plantlet growth divert energy and nutrients away from sexual reproduction, alter plant architecture, and are unnecessary and unwanted in fruit-bearing plants (Heide et al., 2013; Hytönen and Kurokura, 2020; Sønsteby et al., 2021; Voth and Bringhurst, 1990). The runners that emerge in fruit production fields are typically removed by hand to maximize fruit yield, whereas flowers that emerge in bare-root plant (nursery) production fields are commonly removed by hand to maximize daughter plant (clone) yield. The ideal cultivar for a clone-propagation production system produces economically viable clone yields in nursery production and an absolute bare minimum of runners in fruit production. That ideal appears to be difficult to achieve in practice because clone and fruit yield are negatively genetically correlated, although that has not been unequivocally substantiated, partly because runners are normally trimmed in fruit-bearing plants to negate the pleiotropic effect of runner growth on fruit yield.

The ideal cultivars for seed-propagated production systems are runnerless. Although runnerless phenotypes have long been known in the octoploid, the genetic determinants of runnerless phenotypes are unknown and breeding for runnerless and reduced runnering phenotypes has scarcely been documented (Darrow, 1929, 1966).

Little is known about the genetic regulation of asexual reproduction or the heritability of runnering or clone yield variation within short-day and day-neutral populations of octoploid strawberry, apart from the pronounced and well documented pleiotropic effects of the *PERPETUAL FLOWERING* (*PF*) locus on sexual and asexual reproduction (Ahmadi et al., 1990; Bringhurst and Voth, 1976, 1980; Bringhurst et al., 1989; Castro et al., 2015; Gaston et al., 2013; Hossain et al., 2019). Those effects are apparent and exposed when progeny of crosses between short-day (*pfpf*) and day-neutral (*PF_*) parents are grown under long days, typically 13 hours or more (Gaston et al., 2013).

The day-neutral cultivars developed at UC Davis and many others around the globe are descendants of a hybrid between the short-day cultivar ‘Shasta’ and ‘Wasatch’, an ecotype of *F. virginiana* subsp. *glauca* (Bringhurst and Voth, 1980; Ahmadi et al., 1990). That famous ecotype, collected from the Wasatch Mountains of Utah in 1953 by Royce S. Bringhurst, became the backbone of the UC Davis day-neutral breeding program (Bringhurst and Voth, 1976; Bringhurst et al., 1989; Ahmadi et al., 1990; Pincot et al., 2021; Hancock et al., 2008, 2018; Feldmann et al., 2024b).

Bringhurst et al. (1989) and Ahmadi et al. (1990) showed that day-neutral descendants of the original hybrid (‘Shasta’ × ‘Wasatch’) inherited a dominant *PF* allele from ‘Wasatch’ that is necessary for flowering under long days. The ’Wasatch’ *PF* allele, which has been indispensable and widely used in the development of perpetual flowering (day-neutral) cultivars, doubled fruit yields and reshaped strawberry production in California (Hancock, 2006; Hancock et al., 2018; Feldmann et al., 2024b).

The manipulation of flowering phenotypes has been straightforward since the ’Wasatch’ *PF* allele was first introduced into short-day cultivars (Bringhurst and Voth, 1980). The earliest day-neutral cultivars to emerge were ‘Aptos’, ‘Brighton’, and ‘Hecker’ (Bringhurst and Voth, 1980). While those cultivars were groundbreaking, they still harbored unfavorable wild parent alleles and were phenotypically flawed. As the unfavorable alleles and flaws disappeared in their descendants over the next two decades, starting with ’Selva’, the production of DN cultivars progressively expanded until the DN segment dominated production in California and the US (Feldmann et al., 2024b). The ’Wasatch’ *PF* allele simultaneously spread around the globe through the dissemination of UC Davis DN cultivars, which have been widely used in cultivar development (Hancock, 2006; Pincot et al., 2021; Hardigan et al., 2021; Hancock et al., 2018; Feldmann et al., 2024b). Those DN cultivars later spawned the development of so-called extreme day-neutral or summer-plant cultivars that further widened planting and production windows, as exemplified by ’UC Eclipse’ (Knapp et al., 2023).

Although SD and DN plants asexually reproduce under a wide range of conditions, runner and daughter plant growth generally appear to be weaker in *PF_* than *pfpf* individuals under the long days and high temperatures of summer (Gaston et al., 2013). Gaston et al. (2013) genetically mapped a quantitative trait locus (QTL) for runnering (*PFRU*) that co-located with the *PF* locus. They showed that phenotypic differences for runnering were exposed under long days and likely caused by the pleiotropic effects of the *PF* locus. The *PF* locus was originally identified from the Mendelian distributions of long-day flowering habit phenotypes (Ahmadi et al., 1990; Bringhurst and Voth, 1976, 1980; Bringhurst et al., 1989), whereas the *PFRU* QTL was identified from the continuous distributions of runnering and inflorescence count phenotypes (Gaston et al., 2013).

The effects of *PF* alleles are photoperiod and temperature dependent (Bringhurst and Voth, 1976; Gaston et al., 2013; Heide et al., 2013; Hytönen and Kurokura, 2020; Koskela et al., 2016; Sønsteby and Heide, 2006); nevertheless, across the normal range of temperatures encountered in environments that dominate fruit production, *PFPF* and *PFpf* plants flower continuously, whereas *pfpf* plants cease flowering in the late spring and early summer as temperatures increase and daylengths approach the summer solstice (Ahmadi et al., 1990; Bringhurst and Voth, 1976; Bringhurst et al., 1989; Gaston et al., 2013; Heide et al., 2013; Hytönen and Kurokura, 2020). The higher temperatures over the long days of summer typically stimulate runnering in short-day and day-neutral plants (Sønsteby and Heide, 2006). The photoperiod-dependent inhibition of flowering by the recessive (wildtype) allele in short-day (*pfpf*) plants causes a discrete shift from mixed sexual and asexual reproduction to asexual reproduction only, whereas the dominant (mutant) allele (*PF*) knocks out the photoperiod-dependent inhibition of flowering of the wildtype (Ahmadi et al., 1990; Bringhurst and Voth, 1976; Bringhurst et al., 1989; Gaston et al., 2013). Day-neutral (*PF_*) plants flower continuously, exhibit mixed sexual and asexual reproduction over much of their life cycle, and lack the discrete photoperiod-dependent shift from mixed sexual and asexual reproduction to asexual reproduction only (Sønsteby and Heide, 2006). Sønsteby and Heide (2006) stressed that DN strawberries are obligatory long-day plants at high temperature (27°C), quantitative long-day plants at intermediate temperature, and truly day-neutrals only at cool (9°C) temperature. The term day-neutral, however, has become deeply ingrained in the technical lexicon and commerce to describe perpetual flowering (*PF*_) plants, and is used throughout this paper for simplicity, partly because *PF*_ cultivars experience and respond to a wide range of temperatures over an entire growing season in climates where strawberries are produced on a large scale, e.g., coastal California.

While the phenotypic effects of the *PF* locus have been widely discussed (Ahmadi et al., 1990; Bringhurst and Voth, 1976, 1980; Castro et al., 2015; Cockerton et al., 2023; Gaston et al., 2013; Koskela et al., 2016; Lewers et al., 2019; Perrotte et al., 2016; Saiga et al., 2022; Salinas et al., 2017; Verma et al., 2017), the genetic determinants of variation for runnering and clone yield within *PF_* and *pfpf* populations have not. Here we show that the strength of runner suppression is highly variable among *PF_* plants under long days, that runnering is highly variable within populations of short-day (*pfpf*) and day-neutral (*PF_*) plants where the confounding effects of the *PF* locus are absent, and that variation for runnering in short-day and day-neutral plants spans the entire range, from runnerless to prolific and unrestrained. The genetic determinants of that phenotypic variation are unknown in the octoploid; however, clues have emerged from genetic studies in woodland strawberry (*F. vesca*), the diploid donor of the A-genome of the octoploid species (Shulaev et al., 2011; Caruana et al., 2018; Edger et al., 2019; Session and Rokhsar, 2023).

The *PF* locus has been the target of several genetic mapping studies that we show narrowed the location of the *PF* locus down to a 1.0 Mb linkage disequilibrium (LD) block on chromosome 4B (Castro et al., 2015; Cockerton et al., 2023; Gaston et al., 2013; Lewers et al., 2019; Perrotte et al., 2016; Saiga et al., 2022; Salinas et al., 2017; Verma et al., 2017). Those studies left several questions unanswered (reviewed by Hytönen and Kurokura 2020). Importantly, the gene underlying the *PF* locus has not yet been identified, and DNA markers previously developed for marker-assisted selection (MAS) of *PF* alleles are not in complete LD with the *PF* locus and do not always accurately predict *PF* genotypes (Castro et al., 2015; Gaston et al., 2013; Koskela et al., 2016; Lewers et al., 2019; Salinas et al., 2017).

We undertook the studies described here to develop a deeper understanding of the genetics of asexual reproduction and explore genomic prediction as a solution to the problem of accurately predicting runnering phenotypes in octoploid strawberry populations that are either fixed or segregating for *PF* alleles. To accurately predict runnering phenotypes and understand the genetic basis of quantitative variation for runnering in octoploid strawberry, we knew that we needed to correct for the pleiotropic effects of the *PF* locus. Our studies were motivated by the prospect of developing highly productive runnerless cultivars for seed-propagated and near-runnerless cultivars for clone-propagated production systems that strike the perfect balance between sexual and asexual reproduction, within the physiological and developmental limits imposed by the biology of the underlying causal genes, including *PF* (Caruana et al., 2018; Gaston et al., 2013; Koskela et al., 2016; Mouhu et al., 2013; Sønsteby and Heide, 2006).

## 3 MATERIAL AND METHODS

### 3.1 Plant material

The plant materials used in our studies included four seed-propagated F_2_ populations, nine seed-propagated full-sib families, and 932 clonally propagated individuals maintained in the University of California, Davis germplasm collection. The latter spanned the range of runnering phenotypes observed in strawberry, and were selected to construct a strawberry diversity panel (SDP) for genome-wide association and genomic prediction studies. The SDP included 15 *F. chiloensis* ecotypes, 24 *F. virginiana* ecotypes, and 893 *F.* × *ananassa* individuals (Supplemental File S1). The *F.* × *ananassa* individuals included 71 University of California (UC) and 46 non-UC cultivars developed between 1775 and 2017 and 777 hybrid (F_1_) individuals developed between 1937 and 2021. The origin year, taxa, and pedigrees of these individuals are documented in Supplemental File S1.

We developed nine full-sib families by crossing elite SD and DN parents with runner scores ranging from 2.2 to 4.0 (Table 1). The parents included three cultivars released by UCD in 2023 (‘UC Surfline’, ‘UC Monarch, and ‘UC Keystone’) and 11 additional elite F_1_ hybrid individuals selected from full-sib families developed between 2017 and 2019.

**TABLE 1.**
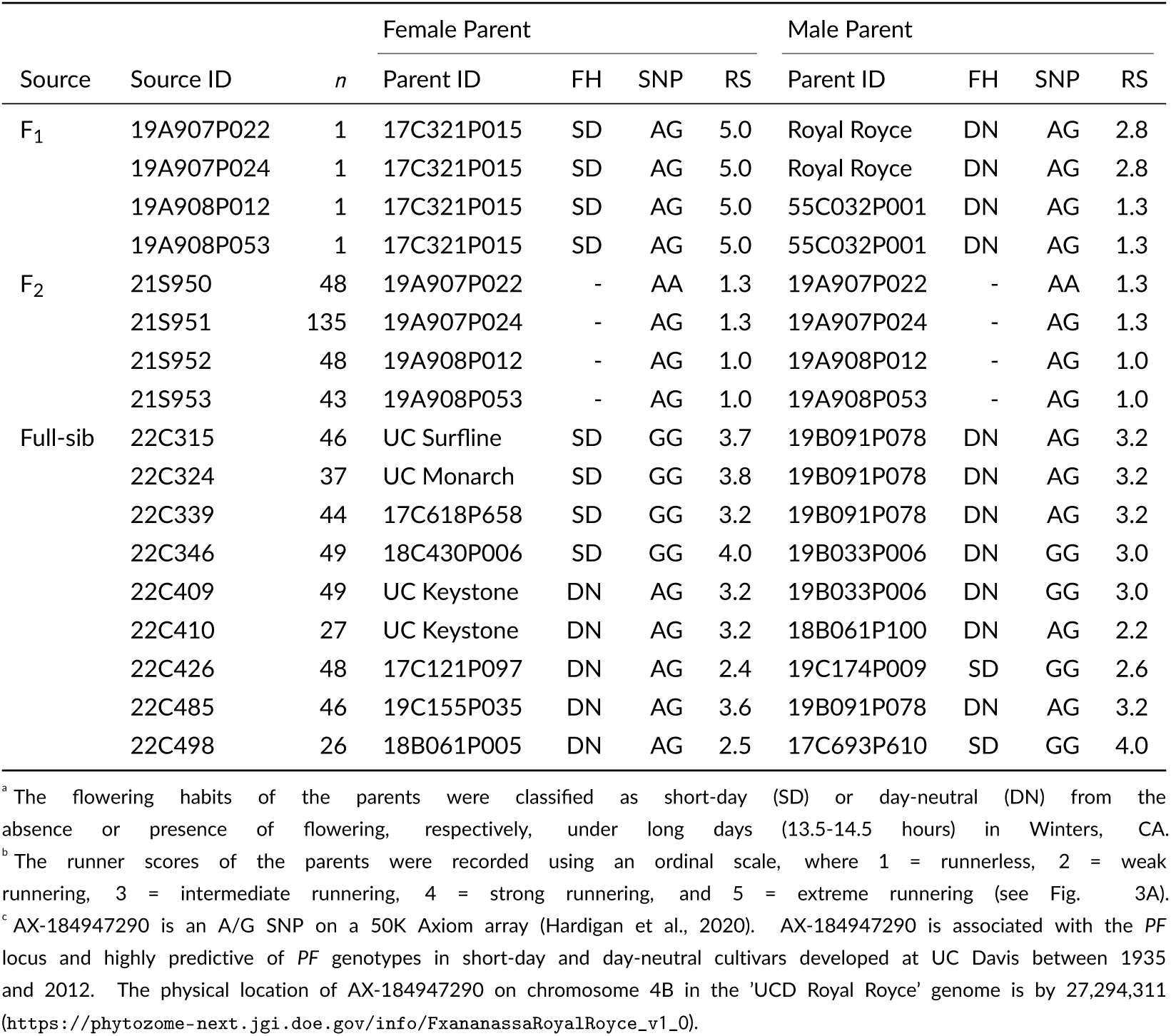
Flowering habit (FH)^a^ and runner score (RS)^b^ phenotypes and A/G SNP marker (AX-184947290) genotypes^c^ of the parents of F_1_ individuals and F_2_ and full-sib families.

F_2_ populations (21S950, 21S951, 21S952, 21S953) were developed by self-pollinating four runnerless to near-runnerless F_1_ individuals (19A907P022, 19A907P024, 19A908P012, and 19A908P053) developed from crosses between parents with contrasting runnering phenotypes (Table 1). The crosses were 17C321P015 × ’Royal Royce’ and 17C321P015 × 55C032P001. We selected two runnerless F_1_ individuals from each full-sib family: 19A907P022 and 19A907P024 from 17C321P015 × ’Royal Royce’ and 19A908P012 and 19A908P053 from 17C321P015 × 55C032P001 (Table 1). The selected individuals were manually self-pollinated over the winter of 2020-21 to produce seeds of two 17C321P015 × ’Royal Royce’ F_2_ populations (21S950 and 21S951) and two 17C321P015 × 55C032P001 F_2_ populations (21S952 and 21S953).

17C321P015 is an SD individual (*pfpf*) with an aggressive runnering phenotype (RS = 5.0) developed from a cross between ‘Cabrillo’ and PI551727. PI551727 (‘CA 1234 Mendocino’) is an SD *F. chiloensis* subsp. *lucida* ecotype with an aggressive runnering phenotype (RS = 5.0) typical of ecotypes of this species. ‘Cabrillo’ is a DN cultivar (*PFpf*) with an intermediate runnering phenotype (RS = 3.0) typical of modern day-neutral cultivars. ‘Royal Royce’ is a DN cultivar (*PFpf*) with slightly weaker runnering (RS = 2.8) than ‘Cabrillo’. 55C032P001 is a runnerless (RS = 1.3), day-neutral individual developed by Royce S. Bringhurst in 1955 from a cross between the SD cultivar ‘Shasta’ (*pfpf*; RS = 3.0) and ‘Wasatch’, an *F. virginiana* subsp. *glauca* ecotype. ’Wasatch’ is the source of the dominant *PF* allele that Bringhurst and Voth (1980) introgressed into short-day genetic backgrounds, starting with ’Shasta’ (Ahmadi et al., 1990).

### 3.2 Field experiments and phenotyping

SDP individuals were grown and clonally propagated annually by rooting plantlets in a field nursery in Winters, CA over several years (2017-2023, excluding 2018). Our study location was the same as that originally used by (Ahmadi et al., 1990) to discover the *PERPETUAL FLOWERING* locus. The annual propagation cycle started with planting two or three bare-root plants/individual in the field in April and ended with the harvest of rooted plantlets (bare-root plants) in December or January. The row spacing was 1.0 m with 0.3 m between plots. Runners were trimmed, if necessary, to prevent cross-contamination between plots. The bare-root plants were dug from the field in December or January of each year, cleaned, trimmed, and stored at 4°for 6-10 weeks before being replanted to repeat the annual propagation cycle.

Over the course of nursery propagation cycle each year, SDP individuals were phenotyped for photoperiod-dependent flowering habit and scored for runnering. The flowering habits of individuals were classified as short-day (SD) or day-neutral (DN) by the absence or presence of flowering, respectively, under long daylengths (13.5-14.5 hours) in late spring and early fall. SDP individuals were phenotyped for runnering in August of each year using ordinal runner scores on a 1-5 scale, where 1 = runnerless, 2 = weak runnering, 3 = intermediate runnering, 4 = strong runnering, and 5 = extreme runnering (see Fig. 3). The runner scores compiled in Supplemental File S1 are estimated marginal means (EMMs) calculated from phenotypes observed over multiple years in Winters, CA. The number of observations per individual ranged from 1-6 (the harmonic mean number of observations per individual was 1.92).

We grew approximately 100 seed-propagated individuals from the 17C321P015 × ’Royal Royce’ and 17C321P015 × 55C032P001 full-sib families in a Winters, CA field experiment in 2018-19. Seedlings of those individuals were grown in a shade house over the summer of 2018, transplanted to the field in September 2018, and phenotyped for runnering in June 2019. Two runnerless individuals were identified and selected within each full-sib family. Those F_1_ individuals were transplanted from the field to a greenhouse in July 2019 and self-pollinated over the winter of 2019-20 to create four F_2_ populations (Table 1).

F_2_ progeny (274 individuals from four families) were phenotyped for runner count in a Davis, CA field experiment in 2021-22 (Table 1). F_2_ seeds were germinated June 2021, seedlings were transplanted to the field in September 2021, and plants were phenotyped by counting primary, secondary, and tertiary runners on 26 March, 9 and 22 April, 6 and 20 May, and 3 and 16 June 2022. The photoperiods on those dates were 12.3, 13.0, 13.3, 14.1, 14.3, 14.4, and 14.5 hours, respectively. The photoperiod on the summer solstice (June 21, 2022) was 14.5 hours.

Full-sib progeny (372 individuals from nine families) were phenotyped for runner score and count in a Davis, CA field experiment in 2023-24. Full-sib seeds were germinated June 2023, seedlings were transplanted to the field in September 2023, and plants were phenotyped by counting inflorescence number and primary runner number on 1 May, 1 June, and 1 July 2024. The photoperiods on those dates were 13.5, 14.4, and 14.5 hr, respectively. Full-sib progeny were scored for runnering as described for the SDP population on 1 July 2024.

### 3.3 Genotyping

Freshly emerged leaves were harvested from SDP, F_2_, and full-sib individuals, lyophilized, and powdered for DNA isolation using the E-Z 96 Plant DNA Kit (Omega Bio-Tek, Norcross, GA, USA) as described by Knapp et al. (2024). DNA samples of these individuals were genotyped using 50K or 850K Axiom^Tm^ SNP genotyping arrays designed by (Hardigan et al., 2020) and developed in collaboration with ThermoFisher Affymetrix Expert Design Program (https://www.thermofisher.com/order/catalog/product/551041. The probe DNA sequences for SNPs on the 50K and 850K Axiom^Tm^ arrays used in our studies were physically anchored to the haplotype-phased ‘UCD Royal Royce’ genome *in silico* (FaRR1; https://phytozome-next.jgi.doe.gov/info/FxananassaRoyalRoyce_v1_0) and unphased ’Camarosa’ genome *in silico* (FaCA1; Edger et al. (2019); https://phytozome-next.jgi.doe.gov/info/ Fxananassa_v1_0_a1). The physical positions of SNPs in the ’UCD Royal Royce’ genome are reported throughout this paper. DNA sequences for the SNP probes and physical coordinates are compiled in Supplemental File S2 for both genomes using the chromosome nomenclature described by Hardigan et al. (2020). That nomenclature is cross-referenced to other linkage group and chromosome nomenclatures in Supplemental File S3. Homoeologous chromosomes in the ’UCD Royal Royce’ genome were numbered 1-7 and oriented according to the rules used for *F. vesca*, the A-genome ancestor (Shulaev et al., 2011; Edger et al., 2018). As previously described (Hardigan et al., 2020), that was feasible because of synteny among homoeologous A, B, C, and D genome chromosomes. DNA samples that passed quality and quantity control standards were genotyped on a GeneTitan HT Microarray System by ThermoFisher Scientific (Santa Clara, CA). SNP genotypes were called using the Affymetrix Axiom Analysis Suite (v1.1.1.66, Affymetrix, Santa Clara, CA).

SNP genotypes were curated and filtered for genome-wide association study (GWAS) and other analyses by retaining those with call rates ≥ 89% and *in silico* assigned physical positions corroborated by *de novo* genetic mapping (the latter are identified in Supplemental File S2). The genomic relationship matrix was estimated among SDP, F_2_, and full-sib individuals using the *rrBLUP::A.mat()* function in the R package ‘rrBLUP’ with a minor allele frequency cutoff of 0.05 (min.MAF = 0.05), a maximum missing data cutoff of 0.8 (max.missing = 0.8), and imputation of missing data (return.imputed = TRUE) (Pincot et al., 2020). After data processing, 41,794 SNPs were retained for analyses of F_2_ and full-sib populations, whereas 42,903 SNP were retained for analyses of the SDP population. We genotyped 194 SDP individuals with the 850K Axiom^Tm^ SNP array (Hardigan et al., 2020). After data processing, 799,895 were retained for analyses of those individuals, whereas 625,039 were retained for analyses of the subset of 66 UC individuals (Supplemental File S2).

Genetic relationships among SDP, F_2_, and full-sib individuals were visualized by constructing a neighbor-joining tree using the *nj()* function in the ’ape’ R package (Paradis et al., 2004) and drawing the tree using the R package ’ggtree’ (Yu et al., 2017). The Euclidean distance matrix for this analyses was estimated from 49,483 array-genotyped SNPs using the *dist()* function in the R ’stats’ package (R Core Team, 2013).

### 3.4 Statistical analyses

The runner scores observed among SDP individuals were analyzed using the *l mer* () linear mixed model (LMM) function in the R package ’lme4’ (Bates et al. 2015; https://cran.r-project.org/web/packages/lme4/index.html),

The LMM for that analysis was

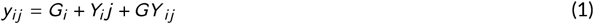

where *y_ij_* is the runner score for the *i* th individual in the *j* th year, *G_i_* is the fixed effect of the *i*th individual, and *Y_j_* is the random effect of the *j* th year, and *GY_ij_* is the random effect of the interaction between individual and year. A single runner score was recorded each year, the composition of the germplasm collection changed year to year, and the number of years per individuals ranged from one to six. The harmonic mean number of replications per individual was 1.92. Estimated marginal means (EMMs) for runner score were estimated using the R package ’emmeans’ (Lenth, 2019).

The individual, year, and individual × year variance components and broad-sense heritability on a clone-mean basis (*H* ^2^ = *V_G_* (*V_G_* + *V_GY_* /*r*) were estimated for runner score using REML in the R package ’lme4’ (Bates et al. 2015, where *V_G_* is the REML estimate of the individual variance component, *V_GY_* is the REML estimate of the individual × year variance component, and *h* = 1.92 is the harmonic mean of the number of observations per individual, calculated using *pysch::harmonic.mean()* (Revelle, 2021).

Genomic-estimated narrow-sense heritability (*h*^2^) was estimated for the SDP, F_2_, and full-sib populations using the *rrBLUP:: mixed.solve()* function, where *h*^2^ = *V_A_*/(*V_A_* + *V_GY_* _)_ /*r*) for the SDP population, *h*^2^ = *V_A_*/(*V_P_* for the F_2_ and full-sib populations, *V_A_* is the genomic-estimated additive genetic variance, and *V_P_* is the phenotypic variance on an individual-plant basis.

Several statistics were estimated for the *PF*-associated SNP AX-184947290 using the LMMs described above. The additive and dominance effects of AX-184947290 were estimated by *a*^ = (*ȳ_AA_* − *ȳ_GG_*)/2 and *d*^^^ = *ȳ_AG_* − (*ȳ_AA_* + *ȳ_GG_*)/2, respectively, where *ȳ_AA_*, *ȳ_AG_*, and *ȳ_GG_* are the respective phenotypic means for AA, AG, and GG SNP genotype. The degree of dominance of AX-184947290 was estimated by |*d*^^^/*a*^| (Falconer, 1996; Walsh, 2001). The percentage of the genetic variance explained by AX-184947290 was estimated by *GV E* = *σ*^ ^2^ /*σ*^ ^2^, where *σ*^ ^2^ is the bias-corrected average semivariance REML estimate of the variance associated with AX-184947290 (Feldmann et al., 2021, 2022). The percentage of the phenotypic variance explained by AX-184947290 was estimated by *PV E* = *σ*^ ^2^ /*σ*^ ^2^ .

### 3.5 Genome-wide association study (GWAS)

Genome-wide association study (GWAS) analyses were applied to the SDP population for flowering habit and runner score, F_2_ population for runner count, and full-sib population for runner score, runner count, and inflorescence count using GEMMA v0.98.1 (Zhou and Stephens, 2012). The runner score EMMs and binary flowering habit classification (SD and DN) were used as input for GWAS of the SDP population. The runner count, inflorescence count, and runner score observations for individual time points were used as input for GWAS of F_2_ and full-sib populations. The GEMMA *‘lmm2’* function and kinship (K) matrix were used to correct for the population structure (Zhou and Stephens, 2012). The *p*-values were adjusted for a false-discovery-rate (FDR) of 0.05 and reported as −*l og*_10_ (FDR p-value) score. Three lead *PF*-associated SNPs (AX-184947290, AX-184219235, and AX-184208850) were used as covariates in GEMMA to search for associations between genetic variants and phenotypes. We tested AX-184947290 and AX-184219235 individually, AX-184947290 and AX-184219235 in combination, and AX-184947290, AX-184219235, and AX-184208850 in combination. Lastly, we used the *GWAS()* function of the ’rrBLUP’ R package for GWAS of SDP individuals genotyped with the 850K SNP array using flowering behavior coded as 0 (short-day) or 1 (day-neutral). The kinship matrix was estimated using the *rrBLUP::A.mat()* function and incorporated in the ’rrBLUP’ analysis to control for population structure.

### 3.6 Haplotype analyses

The linkage phases of 50K array-genotyped SNPs were unknown among individuals in the SDP and F_2_ populations. They were inferred in the latter by genetic mapping and by comparing haplotypes inferred among SDP individuals. We used Beagle 5.4v (Browning et al., 2021) with default parameters to infer the haplotypes of 51 SNPs that were associated with the *PF* locus and had minor allele frequencies (MAFs) ≥ 0.3 in the SDP population. GWAS-estimated SNP effects with FDR-corrected −*l og*_10_ (p-value) > 2 for flowering habit or runner score were selected for phasing. This filtering process yielded 51 SNPs spanning bp 26,529,027 to 29,635,684 on chromosome 4B. The squared correlation coefficient linkage disequilibrium statistic (*R* ^2^) was estimated among the 51 SNPs using Beagle 5.4v (Browning et al., 2021).

### 3.7 Genomic prediction analyses

Genomic relationship matrices (G) and genomic-estimated breeding values (GEBVs) were estimated by genomic best linear unbiased prediction (G-BLUP) for runner score in the SDP population, runner count in the F_2_ population, and runner count and and inflorescence count in the full-sib population using the *mixed.solve()* function in the R package rrBLUP (Endelman, 2011; Endelman and Jannink, 2012)). GEBVs were estimated with and without incorporating the AX-184947290 SNP as a fixed effect in the G-BLUP analysis. Genomic predictive ability (*r_a_*_^,*ȳ*_) and prediction accuracy (*r_a_*_^,*ȳ*_ /*h*) were estimated for runner score or count within and between strawberry populations by cross-validation, where *a*^ is the GEBV and *ȳ* is the EMM. We used the SDP population (*n* = 932) and subset of UC individuals (*n* = 672) in the SDP population (SDP-UC) for cross-validation. Cross-validation was performed with 1,000 Monte Carlo iterations within the SDP, SDP-UC, and full-sib populations and between the SDP and full-sib and SDP-UC and full-sib populations by randomly splitting individuals into training (80%) and validation (20%) subsets (Burman, 1989; Molinaro et al., 2005; Poland and Rutkoski, 2016).

## 4 RESULTS AND DISCUSSION

### 4.1 Variation for runnering encompasses the entire phenotypic range, from runnerless to prolific

Fig. 1-3 depict the range of stolon (runner) and plantlet (daughter plant) growth phenotypes observed among several thousand seed-propagated full-sib progeny in early-stage selection nurseries and 3,000 clonally propagated short-day and day-neutral individuals observed in the UC strawberry germplasm collection over a period of nine years (2015-2023). That variation was captured by the sample of 932 clonally propagated strawberry diversity panel (SDP) individuals phenotyped for photoperiod-dependent flowering habit and runnering in a low-elevation nursery production environment (Winters, CA; 38.52°N, 121.97°W; 41 m) where photoperiods ranged from 9.5 hr to 14.9 hr over the March to December growing season (Fig. 3; Supplemental File S1). The term “runnering” applied here and throughout our paper, though informal, is commonly used in research and commerce to describe stolon (runner) and plantlet (daughter plant) growth phenotypes.

**FIGURE 1.**
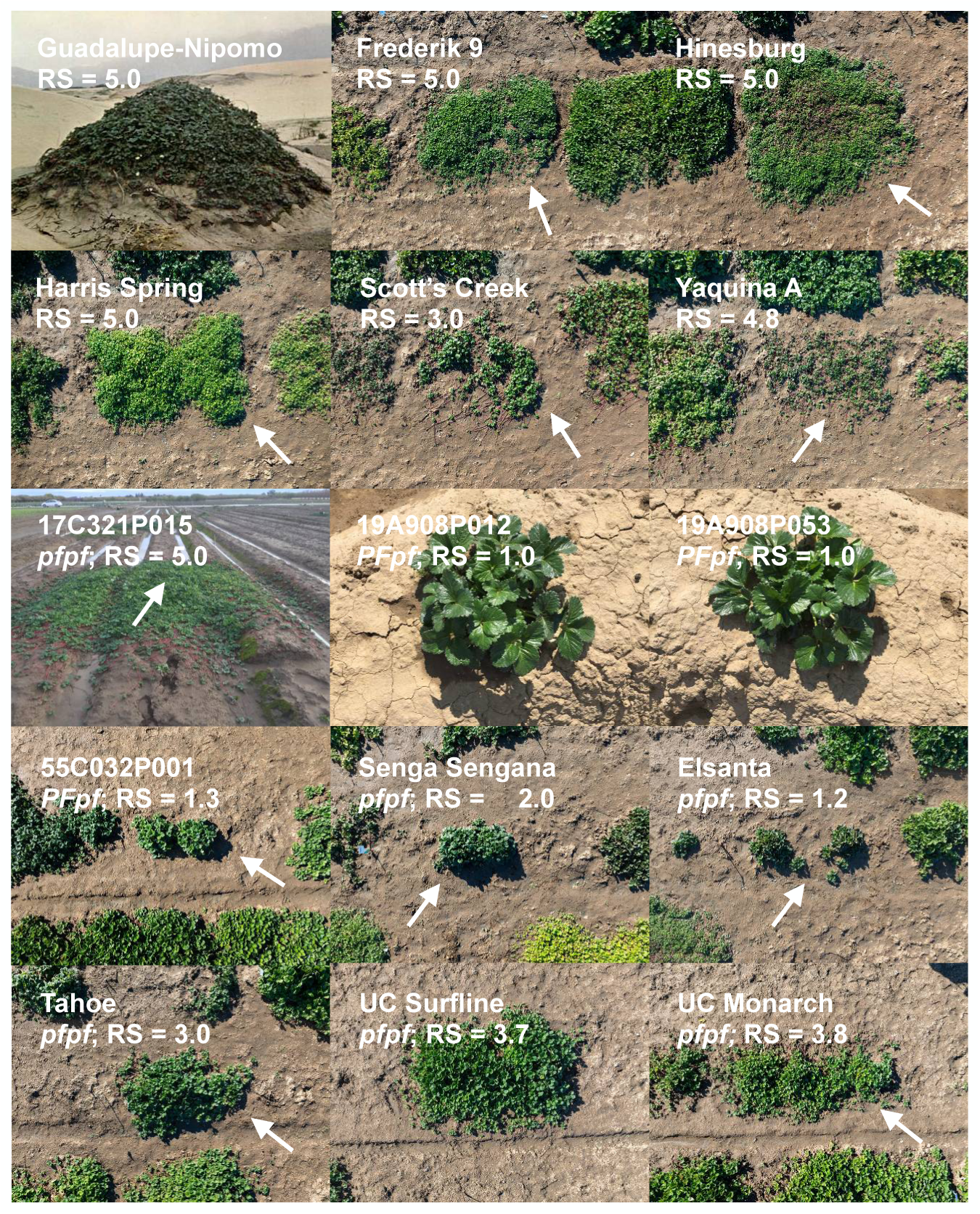
Variation for runnering in octoploid strawberry. Unless otherwise noted, the phenotypes shown in these photographs (shot to scale by Fred Greaves for UC Davis) were observed 11 November 2024, Winters, CA. ’Guadalupe-Nipomo’, an *F. chiloensis* subsp. *lucida* ecotype carpeting a sand dune near Oso Flaco Beach, CA (August 1963). ’Frederik 9’ and ’Hinesburg’ are *F. virginiana* subsp. *virginiana* ecotypes. ’Harris Spring’ and ’Scott’s Creek’ are *F. virginiana* subsp. *platypetala* ecotypes. ’Yaquina A’ is an *F. chiloensis* subsp. *pacifica* ecotype. 19A908P012 and 19A908P053 are runnerless 17C321P105 × 55C032P001 hybrids (July 2020, Winters, CA). 17C321P015 is a hybrid between ’Cabrillo’ and PI551727, an *F. chiloensis* subsp. *lucida* ecotype (December 2017, Winters, CA). 55C032P001 is a runnerless day-neutral hybrid. ’Senga Sengana’, ’Elsanta’, ’Tahoe’, ’UC Surfline’, and ’UC Monarch’ are short-day cultivars. *PF* genotypes were predicted from the segregation of *PF* in full-sib families and by using a SNP marker (AX-184947290) associated with the *PF* locus. RS is the estimated marginal mean for runner score over years. Arrows identify the labeled individuals where unrelated individuals appear in growth boundaries.

**FIGURE 2.**
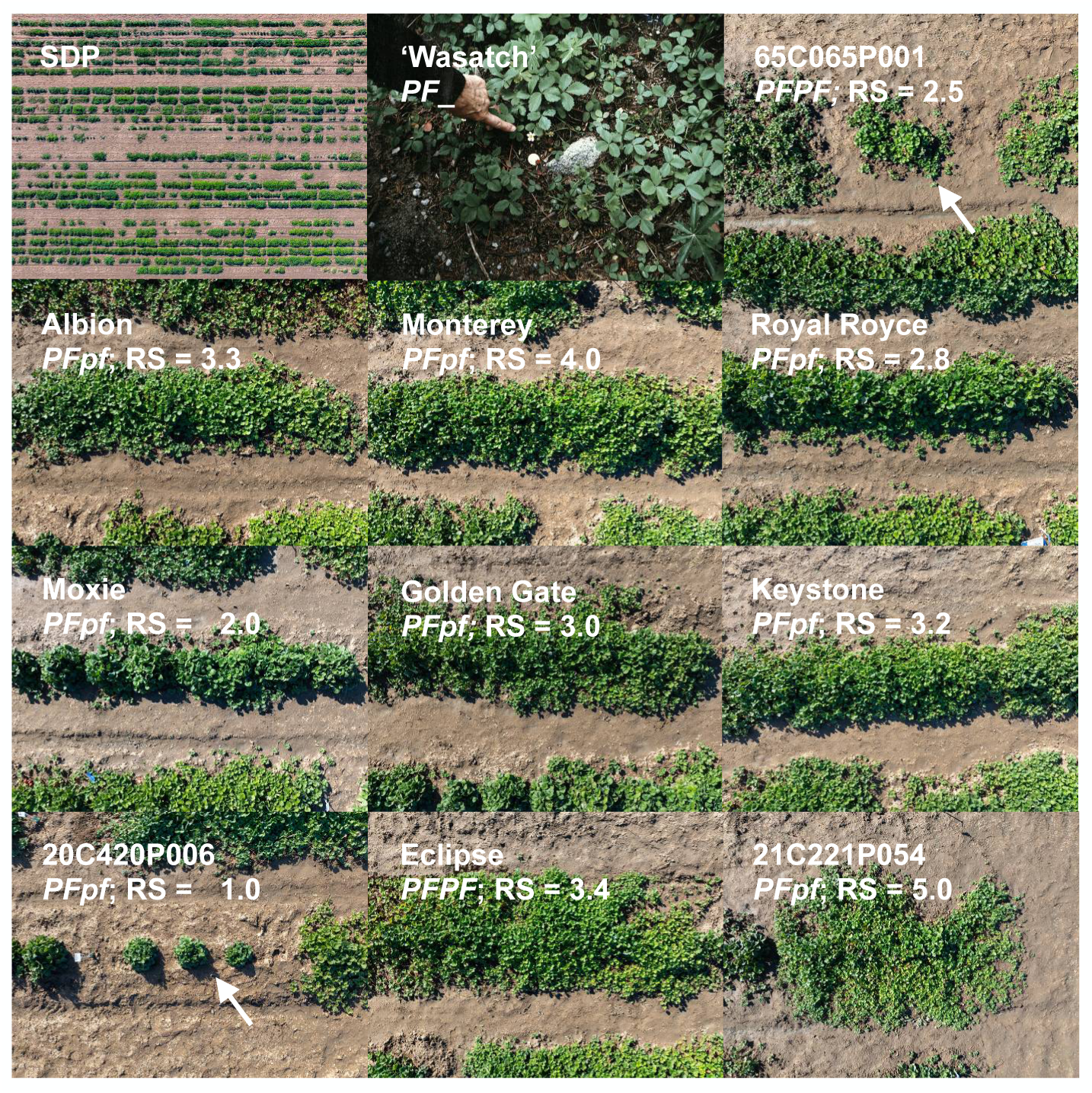
Variation for runnering among day-neutral (*PFPF* and *PFpf*) individuals known to be descendants of ‘Wasatch’, the *F. virginiana* subsp. *glauca* donor of the *PF* allele. The ’Wasatch’ photograph shows the original *PF* donor plant collected July, 1953 from the Wasatch Mountains of Utah (reproduced from an original photograph in the Royce S. Bringhurst Special Collection (UUS COLL MSS 515), Merrill-Cazier Library, Utah State University; https://library.usu.edu/archives/). The phenotypes shown in the other photographs (shot to scale by Fred Greaves for UC Davis) were observed 11 November 2024 in Winters, CA. *PF* genotypes were predicted using a SNP marker (AX-184947290) associated with the *PF* locus. RS is the estimated marginal mean for runner score over years. Arrows identify the labeled individuals where unrelated individuals appear in growth boundaries. The aerial photograph (upper left corner) displays a subset of the SDP and other individuals preserved in the clonal germplasm collection at UC Davis.

**FIGURE 3.**
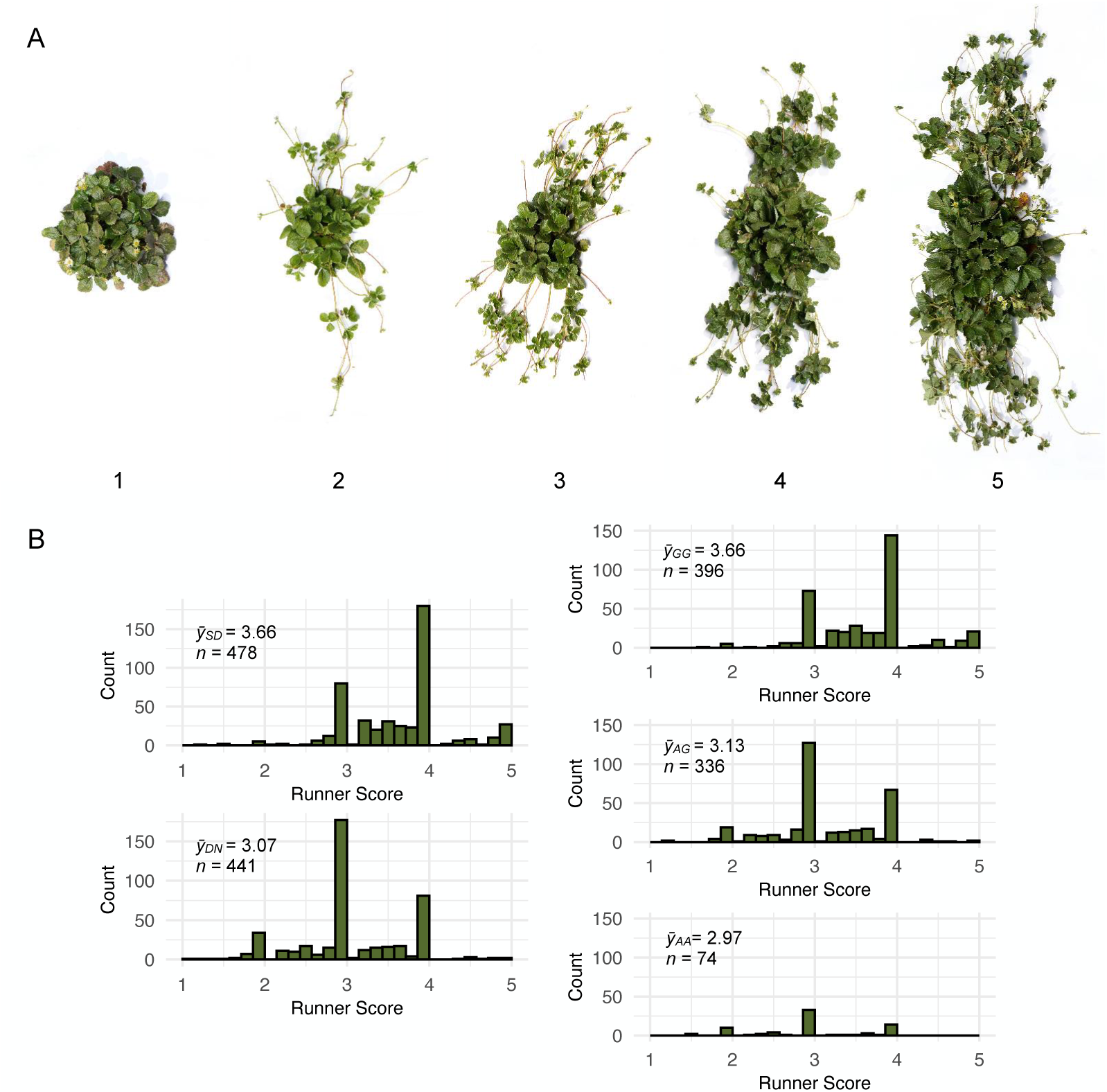
Runner score distributions of strawberry diversity panel individuals classified as short-day (SD = *pfpf*; *n* = 479) or day-neutral (DN = *PF*_; *n* = 441) over six growing seasons in Winters, CA. (A) Archetypal phenotypes of individuals scored for runnering on an ordinal scale, where 1 = runnerless, 2 = weak runnering, 3 = intermediate runnering, 4 = strong runnering, and 5 = extreme runnering (shot to scale by Fred Greaves for UC Davis, July 2024, Winters, CA). (B) Histograms were plotted using the phenotypic means (EMMs = *ȳ*) for runner scores over years (the harmonic mean number of observations per individual was 1.92). The *ȳ* distributions of SDP individuals classified as SD or DN are shown in the left column with their group means (*ȳ_SD_* and *ȳ_DN_*). The *ȳ* distributions of SDP individuals with GG, AG, and AA AX-184947290 SNP genotypes are shown in the right column of histograms with their genotypic means (*ȳ_GG_*, *ȳ_AG_*, and *ȳ_AA_*).

The SDP included 15 *F. chiloensis* ecotypes, 24 *F. virginiana* ecotypes, and 893 *F.* × *ananassa* individuals with runner and daughter plant growth (runnering) phenotypes spanning the range found in the octoploid, from runnerless to prolific and unrestrained (Fig. 1-3; Supplemental Files S1). Genetic relationships among these individuals are depicted in Fig. 4, along with individuals from two exotic × exotic (17C321P015 × 55C032P001) F_2_ families, two elite × exotic (17C321P015 × ’UCD Royal Royce’) F_2_ families, and nine elite × elite full-sib families that were phenotyped for runnering in our study (Table 1; Supplemental Files S4-S5). The nine full-sib families were developed from crosses among 13 elite UC parents (Table 1).

**FIGURE 4.**
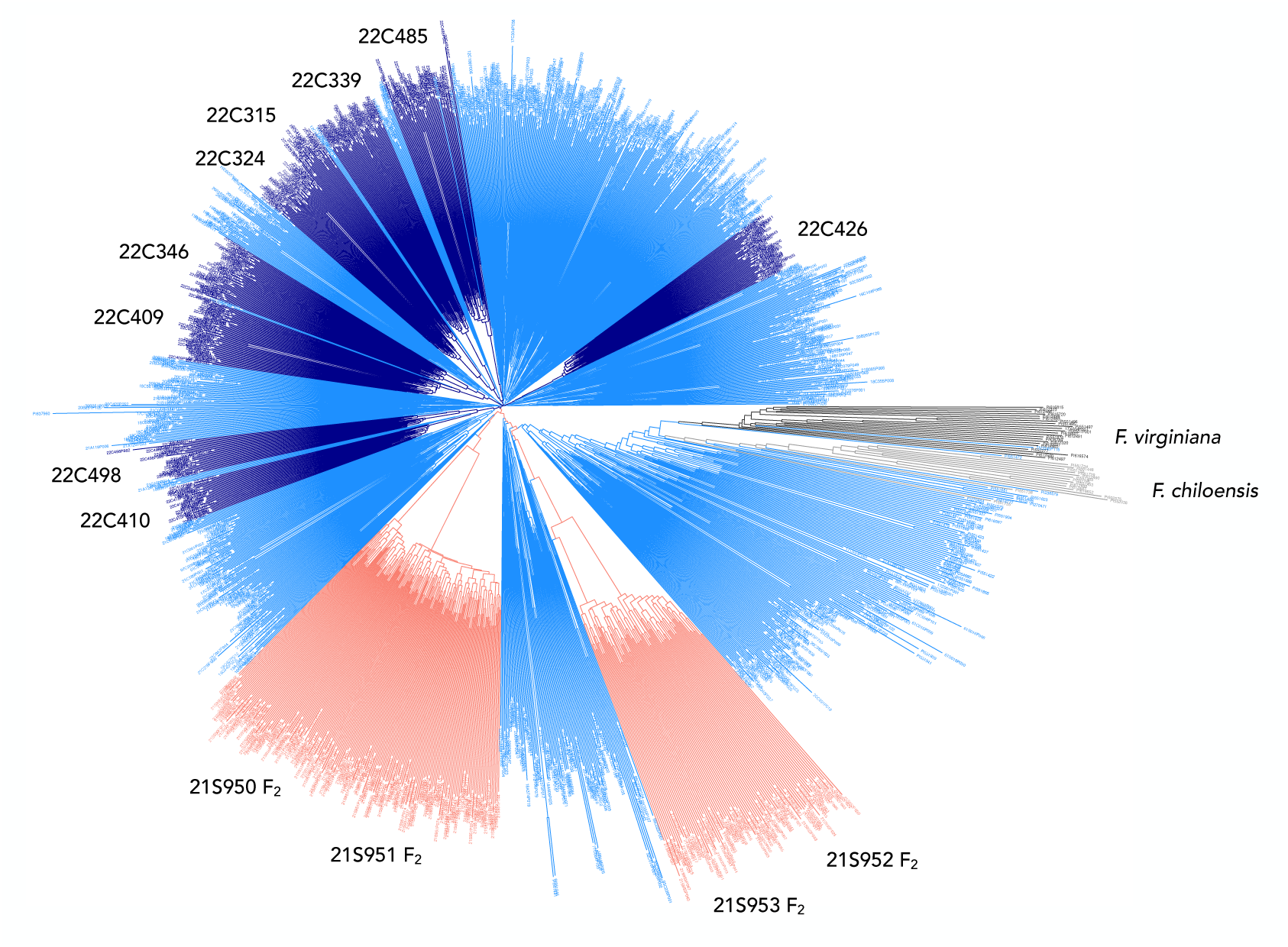
Genetic relationships among 932 strawberry diversity panel, 270 F_2_, and 372 full-sib individuals genotyped with an Axiom 50K SNP array. Genetic relationships were estimated from 49,483 SNPs. Clades identified by 21C950 and 21C951 are 17C321P015 × ’UCD Royal Royce’ F_2_ families. Clades identified by 21C952 and 21C953 are 17C321P015 × 55C032P001 F_2_ families. Clades identified by 22C numbers are full-sib families developed from crosses between elite short-day and day-neutral individuals between 2017 and 2019 at UC Davis. The other branches and nodes identify strawberry diversity panel individuals, including *F. chiloensis* and *F. virginiana* ecotypes.

We knew from previous studies that we needed to correct for the photoperiod-dependent pleiotropic effects of the dominant ’Wasatch’ *PF* allele to identify loci affecting runnering phenotypes (Ahmadi et al., 1990; Gaston et al., 2013). SDP individuals were classified as short-day or day-neutral by the presence or absence of flowering under longdays (14.0-14.9 hr), respectively (Fig. 3B; Supplemental File S1). The repeatability of our flowering habit classifications was 99% over six growing seasons (March to December) in Winters, CA (38.52°N). Short-day individuals were inferred to be homozygous recessive (*pfpf*), whereas day-neutral individuals were inferred to be heterozygous or homozygous dominant (*PF*_) for *PF* alleles (Ahmadi et al., 1990; Bringhurst and Voth, 1976; Bringhurst et al., 1989; Gaston et al., 2013; Perrotte et al., 2016).

SDP individuals were phenotyped for runnering in our clonal propagation nursery in Winters, CA over six growing seasons (March to December) using an ordinal runner score (RS) scale, where 1 = runnerless, 2 = weak runnering, 3 = intermediate runnering, 4 = strong runnering, and 5 = extreme runnering (Fig. 3). The number of years of observation ranged from one to six per individuals (the harmonic mean number of years of observation was 1.92). To complement our analyses of full-season phenotypes of clonally propagated SDP individuals, seed-propagated exotic × exotic (17C321P015 × 55C032P001) F_2_, elite × exotic (17C321P015 × ’UCD Royal Royce’) F_2_, and elite × elite full-sib progeny were phenotyped by counting runners under long days (13.0 to 14.5 hr photoperiods in Davis and Winters, CA). The runner count phenotypes of F_2_ individuals were recorded on seven dates with photoperiods ranging from 13.0 to 14.5 hr (Supplemental File S4). Our experience in the F_2_ study guided the phenotyping schedule applied in the full-sib study. Full-sib individuals were phenotyped on three dates (1 May, 1 June, and 1 July) with photoperiods ranging from 13.5 to 14.5 hr (Supplemental File S5). Unless otherwise noted, we are reporting statistics estimated from phenotypes observed on the longest days when phenotypic differences and the temporal effect of the *PF* locus were the greatest.

The individuals in our study populations sampled genetic variation across the entire diversity range, from the most exotic (ecotypes of the wild relatives) to the most elite (modern UC cultivars). Among SDP individuals, 478 were classified as short-day and 441 were classified as day-neutral (Fig. 3B; Supplemental File S1). The *F.* × *ananassa* individuals included 80 short-day cultivars, 39 day-neutral cultivars, 776 UC cultivars and other hybrids developed between 1969 and 2021 (the SDP-UC training population), and 691 UC cultivars and other hybrids developed between 2016 and 2021. The 893 *F.* × *ananassa* individuals selected for the highly admixed SDP population originated from 554 unique pedigrees (full-sib families) among 489 parents (Fig. 4; Supplemental File S1).

The runnering phenotypes shown in Fig. 1-2, with four exceptions (’Guadalupe-Nipomo’, 17C321P015, 19A908P012, and 19A908P053), were observed 11 November 2024 after a full-season of runner growth and clonal propagation in Winters, CA, and were shot to scale to ensure accurate visual comparisons (the mother plants were transplanted March 2024). The runners of those individuals were periodically trimmed to prevent cross-contamination between individuals; hence, the phenotypes shown for wild relatives and other individuals with prolific and unrestrained runnering are less extreme than would have been observed without runner trimming.

Untrimmed phenotypes are shown in Fig. 1 for an *F. chiloensis* subsp. *lucida* ecotype (’Guadalupe-Nipomo’) *in situ* and 17C321P015 *ex situ*, a hybrid between the UC day-neutral cultivar ’Cabrillo’ and an *F. chiloensis* subsp. *lucida* ecotype (PI551727) native to Cape Mendocino, CA. The iconic photograph of the ’Guadalupe-Nipomo’ ecotype, captured by Royce S. Bringhurst near Oso Flaco Beach, CA, illustrates the coastal adaptation and prolific and unrestrained runner growth of this wild relative in a sand dune habitat (Bringhurst et al., 1977). The phenotype shown for 17C321P015 depicts a single season of nursery growth from a single mother plant (identified by the arrow) without runner trimming (Fig. 1). The diameter of the labyrinth of runners and daughter plants emanating from the 17C321P015 mother plant was approximately 3.7 meters at the end of the clonal propagation season (December 2017).

At the opposite extreme are the runnerless to near-runnerless phenotypes of several day-neutral ’Wasatch’ descendants, e.g., 55C032P001 (55.32-1), 20C420P006, and ’UCD Moxie’ (Fig. 1-2). 55C032P001 is a first-generation, day-neutral descendant of ’Shasta’ × ’Wasatch’ and donor of the *PF* allele found in 65C065P601, ’Aptos’, ’Brighton’, and ’Hecker’, the earliest UC day-neutral cultivars (Bringhurst and Voth, 1980; Ahmadi et al., 1990; Pincot et al., 2021). 65C065P601 (65.65-601) is a third-generation day-neutral descendant of ‘Shasta’ × ‘Wasatch’, parent of the DN cultivar ‘Brighton’, and grandparent of the DN cultivar ‘Selva’ (Ahmadi et al., 1990; Bringhurst and Voth, 1980). High-density SNP haplotypes for those individuals facilitated analyses to pinpoint the physical location of the *PF* locus.

### 4.2 The runnering phenotypes of strawberry are highly heritable and stable

Although runnerless and near-runnerless phenotypes have long been known in so-called “ever-bearing” cultivars of octoploid strawberry (Darrow, 1929, 1966), the runnerless phenotype of the 55C032P001 individual and other perpetual flowering descendants of the ’Wasatch’ ecotype have apparently never been documented (Fig. 1-3). The emergence of that transgressive phenotype among ’Shasta’ × ’Wasatch’ progeny must have been unanticipated by Bringhurst and Voth (1980) because both parents are prolific runner producers (the mean runner score for ’Shasta’ was 3.0 in our study). The original ‘Wasatch’ ecotype disappeared from the UC collection long ago (Pincot et al., 2021); however, the ecotype recollected by Hancock et al. (2001) from the original ’Wasatch’ population (BT3; PI612491) was shown to have a prolific and unrestrained runnering phenotype typical of *F. virginiana* subsp. *glauca* (the mean runner score for BT3 was 4.83 in our study).

Several transgressive phenotypes were recovered in our breeding and genetic studies, e.g., the runnerless phenotypes of the F_1_ parents of our 17C321P015 × ’Royal Royce’ F_2_ populations (19A907P022 and 19A907P024) were transgressive (Table 1; Fig. 1-3). The runners scores for both were 1.3, whereas the runner scores of the parents were 2.8 and 5.0. Conversely, the runnerless to near-runnerless phenotypes recovered among 17C321P015 × 55C032P001 progeny was anticipated because the 55C032P001 parent was runnerless (RS = 1.3).

The runnerless phenotype of 55C032P001 has been unaffected by temperature and photoperiod variation over nine field growing seasons (March-December) in Winters, CA (Fig. 1). Similarly, the phenotypes of 20C420P005 and the other runnerless to near-runnerless individuals identified in our studies have been stable and predictable across nursery and fruit production environments (Fig. 1-2; Supplemental File S1). Our estimate of broad-sense heritability on a clone-mean basis (*H* ^2^) was 0.86 for runner score in the SDP population (Table 2). Our estimates of narrow-sense heritability (*h*^2^) for runner score or count in the SDP, F_2_, and full-sib populations were more moderate, ranging from 0.37 to 0.60 (Table 2).

**TABLE 2.** Additive genetic variance (*V_A_*), coefficient of additive genetic variance (*CV_A_*), and heritability of runnering phenotypes in strawberry.

| Population<sup>a</sup> | Trait<sup>b</sup> | *n* | *y*<sub>MIN</sub> | *y*<sub>MAX</sub> | *V*<sub>A</sub> | *CV*<sub>A</sub> | *h*<sup>2</sup> | *H*<sup>2</sup> |
| --- | --- | --- | --- | --- | --- | --- | --- | --- |
| Diversity Panel | Runner Score | 932 | 1 | 5 | 0.3 | 15.9 | 0.60 | 0.86 |
| F<sub>2</sub> Families | Runner Count | 270 | 0 | 104 | 58.5 | 32.7 | 0.37 | - |
| Full-sib Families | Runner Count | 372 | 4 | 98 | 183.9 | 30.7 | 0.53 | - |
|  | Inflorescence Count |  | 0 | 41 | 19.1 | 177.7 | 0.60 | - |

<sup>a</sup> Statistics were estimated from clonal replicates of strawberry diversity panel individuals, unreplicated seed-propagated 17C321P015 × 'Royal Royce' and 17C321P015 × 55C032P001 F<sub>2</sub> progeny, and unreplicated seed-propagated full-sib progeny from nine crosses between elite parents with contrasting runnering phenotypes (Table 1). *n* is the number of individuals in the population. *y*<sub>MIN</sub> and *y*<sub>MAX</sub> are phenotypic minimums and maximums, respectively. Genetic variance among clones of individuals (*V*<sub>G</sub>) and broad-sense heritability on a clone-mean basis ( $H^2 = V_G/V_{\bar{p}}$ ) were estimated for runner score among strawberry diversity panel (SDP) individuals, where *V* <sub>$\bar{p}$</sub> is the phenotypic variance on a clone-mean basis. Genomic additive genetic variance (*V*<sub>A</sub>) and narrow-sense heritability on a clone-mean basis ( $h^2 = V_A/V_{\bar{p}}$ ) were estimated for runner score among SDP individuals. *V*<sub>A</sub> and narrow-sense heritability on a individual-plant basis ( $h^2 = V_A/V_p$ ) were estimated for runner count among F<sub>2</sub> progeny and runner and inflorescence count among full-sib progeny, where *V*<sub>*p*</sub> is the phenotypic variance on an individual plant basis. $CV_A = \sqrt{V_A}/\bar{y} \times 100$ , where $\bar{y}$ is the population mean.
<sup>b</sup> The flowering habits of SNP individuals were classified as short-day or day-neutral from the absence or presence of flowering, respectively, under long days (13.5-14.5 hours) over a period of six years in Winters, CA. The runner scores of SDP individuals were similarly recorded over a period six years in Winters, CA using an ordinal scale, where 1 = runnerless, 2 = weak runnering, 3 = intermediate runnering, 4 = strong runnering, and 5 = extreme runnering (see Fig. 3A). F<sub>2</sub> and full-sib progeny were phenotyped by counting the number of primary runners observed in Davis or Winters, CA, respectively, within one week of the summer solstice (the daylength was 14.5 hours).

### 4.3 Physical mapping uncovered the persistence of LD-dragged DNA among ’Wasatch’ descendants carrying the *PF* allele

We physically mapped the *PF* locus to the lower arm of chromosome 4B using associations between octoploid genome-anchored 50K array SNPs and long-day flowering habit classifications of SDP individuals (Fig. 5A-6A). Statistically significant associations were not observed elsewhere in the genome apart from singletons on chromosomes 4C and 7A that were identified in our initial genome-wide search and subsequently found to be false-positives (Fig. 5A). The lead *PF*-associated SNPs were AX-184219235 (−*l og* 10(*p*) = 35.7; bp 27,206,278) and AX-184947290 (−*l og* 10(*p*) = 34.5; bp 27,294,311).

**FIGURE 5.**
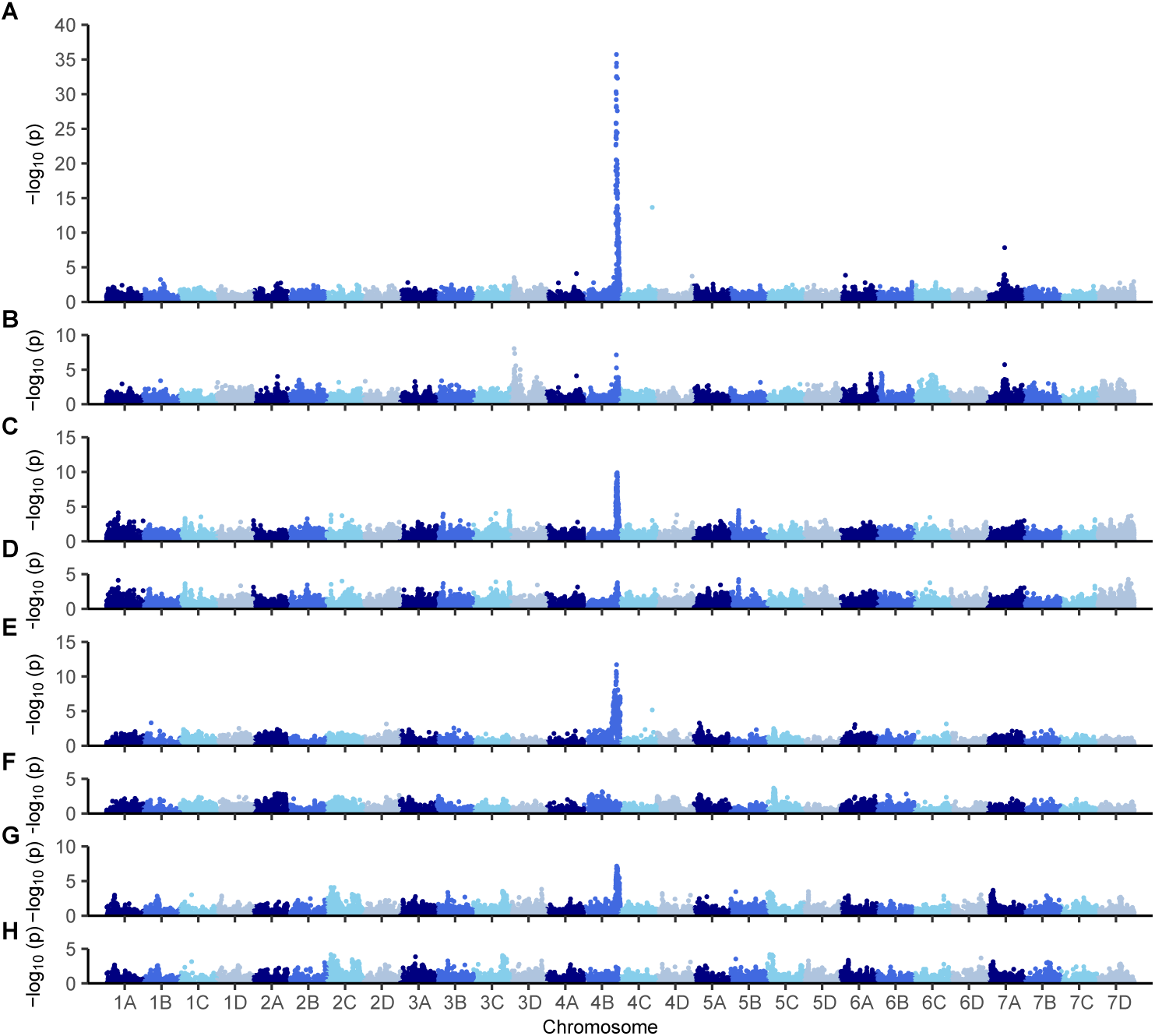
Manhattan plots displaying Axiom 50K array-genotyped SNPs associated with flowering or runnering phenotypes across the octoploid strawberry genome. There are four pairs of plots (A-B, C-D, E-F, and G-H) displaying statistics for GWAS without fixed effects (upper panel in each pair) and where the *PF*-associated SNP AX-184947290 was incorporated as a fixed effect (lower panel in each pair). (A-B) GWAS for flowering habit among 932 strawberry diversity panel individuals. (C-D) GWAS for runner score among 932 strawberry diversity panel individuals. (E-F) GWAS for runner count among 183 17C321P015 × ’UCD Royal Royce’ F_2_ and 87 17C321P015 × 55C032P001 F_2_ progeny. (G-H) GWAS for runner count among 372 full-sib progeny developed from crosses between elite UC parents.

**FIGURE 6.**
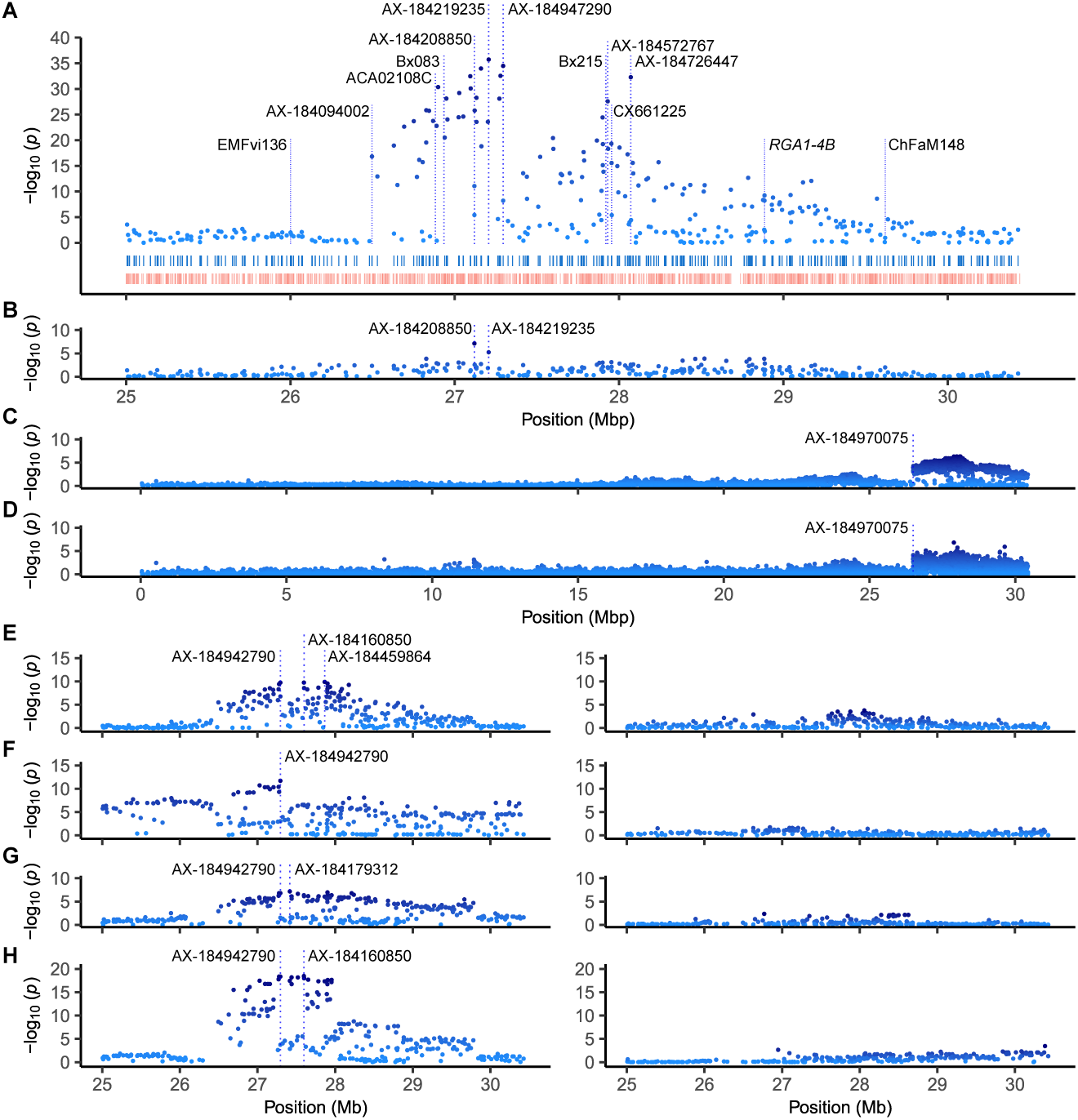
Manhattan plots displaying Axiom 50K or 850K array-genotyped SNPs associated with flowering or runnering phenotypes on chromosome 4B. The physical locations of several array-genotyped SNPs (identified by AX prefixes), previously genetically mapped DNA markers (identified by other labels), and *RGA1-4B* (a *REPRESSOR OF GA1* homoeolog) are shown in the ’UCD Royal Royce’ genome. The Manhattan plots shown in B and the right hand column of E-H display associations identified when AX-184947290 was used as a fixed effect. (A) SNPs associated with flowering habit observed among 932 strawberry diversity panel (SDP) individuals. The upper rug plot shows the physical positions of 322 Axiom 50K array SNPs, whereas the lower rug plot shows the positions of 905 annotated genes spanning Mb 25.0-30.4 in the ’UCD Royal Royce’ genome. (B) SNPs associated with flowering habit among 932 SDP individuals identified by GWAS using AX-184947290 as a fixed effect. (C) SNPs associated with flowering habit among 66 UC individuals genotyped with the 850K array. (D) SNPs associated with flowering habit among 194 SDP individuals genotyped with the 850K array. (E) SNPs associated with runner score among 932 SDP individuals. (F) SNPs associated with runner count among 183 17C321P015 × ’UCD Royal Royce’ F_2_ and 87 17C321P015 × 55C032P001 F_2_ progeny. (G) SNPs associated with runner count among 372 full-sib progeny developed from crosses between elite UC parents. (H) SNPs associated with inflorescence count among 372 full-sib progeny developed from crosses between elite UC parents.

The pattern of linkage disequilibrium (LD) decay for *PF*-associated SNPs was mostly smooth in the SDP population (Fig. 6A). LD steadily decayed upstream of AX-184947290 to an LD cliff (AX-184094002; bp 26,496,157) and downstream of AX-184947290 to the nearly the end of the chromosome. There was a trough and irregular pattern of LD decay immediately downstream of AX-184947290 (Fig. 6A). Two SNPs downstream of trough were strongly associated with the *PF* locus, AX-184572767 (−*l og* 10(*p*) = 27.69; bp 27,930,755) and AX-184726447 (−*l og* 10(*p*) = 32.28; bp 28,070,877). We discovered that those associations were likely caused by the highly admixed structure of the SDP population (Hou et al., 2021), not by the segregation of a second locus.

Because the effect of the *PF* locus dwarfs and could mask the effects of other loci affecting photoperiod-dependent flowering, GWAS was repeated using *PF*-associated SNPs as fixed effects, specifically AX-184219235 or AX-184947290 alone, AX-184219235 and AX-184947290 in combination, and AX-184219235, AX-184947290, and AX-184208850 in combination (Fig. 5B-6B). These were the three most highly predictive *PF*-associated SNPs for flowering habit in the SDP. When AX-184947290 alone was used as a fixed effect for GWAS, the effects of SNP singletons on chromosomes 4C and 7A, the SNP outliers downstream of AX-184947290 on chromosome 4B (AX-184572767 and AX-184726447), and other *PF*-associated SNPs on chromosome 4B either nearly or completely disappeared (Fig. 5B-6B). The effects of AX-184208850 and AX-184219235 were slightly above the genetic background when AX-184947290 was used as a fixed effect and vice versa for the others (Fig. 6B). Our results suggested that the fixed effects captured haplotypic diversity in the highly admixed SDP population (Hou et al., 2021) and that the pattern of LD decay observed in the SDP population was caused by the segregation of a single gene in strong LD with AX-184947290 (bp 27,294,311).

We genotyped a small sample of short-day and day-neutral UC individuals (*n* = 66) and a slightly larger sample of more diverse short-day and day-neutral individuals (*n* = 194) with an 850K SNP array to graphically genotype chromosome 4B and the ’Wasatch’ introgression using GWAS (Fig. 6C). While GWAS analyses of those small samples were underpowered and insufficient for accurately pinpointing the physical location of *PF*, that analysis saturated the ’Wasatch’ introgression with 2,600 SNPs, clearly identified a stable recombination breakpoint upstream of the *PF* locus (the upper boundary of the ’Wasatch’ introgression), and showed that ’Wasatch’ DNA has persisted and survived selection for perpetual flowering from the recombination breakpoint to the nearly the end of the chromosome since the original hybrid (’Shasta’ × ’Wasatch’) was developed in 1955 (Fig. 6C). The upper boundary of the ’Wasatch’ introgression was marked by an LD cliff where significance dropped to near zero immediately upstream of the 850K array SNP AX-184970075 (bp 26,490,375) (Fig. 6A and C-D). The recombination breakpoint was predicted to reside between AX-184284360 (bp 26,484,790) and AX-184970075 (26,490,375) among individuals genotyped with the 850K SNP array. That interval was inside the interval (bp 26,480,809-26,496,157) where the recombination breakpoint was predicted to reside in individuals genotyped with the 50K SNP array.

The invariance of the upper boundary suggests that the location of crossovers has been highly constrained in ’Wasatch’ descendants that inherited the *PF* allele (Fig 6A and C-D). We observed runs-of-homozygosity and suppressed recombination (a recombination cold spot) immediately upstream of the recombination breakpoint (Fig. 6A and C-D). The LD pattern upstream of the recombination breakpoint could correlate with structural variation between short-day recipients and the ’Wasatch’ donor, pedigree inbreeding caused by backcrossing to closely related short-day recipients, or a combination thereof (Wright and Andolfatto, 2008; Ceballos et al., 2018).

### 4.4 The integration of physically and genetically mapped DNA markers associated with the *PF* locus

Our study facilitated the integration and cross-referencing of findings from earlier genetic mapping studies of the *PF* locus (Sargent et al., 2016; Spigler et al., 2008, 2010; Perrotte et al., 2016; Honjo et al., 2016; Saiga et al., 2022). Historically, that has been challenging because of differences in linkage group and chromosome nomenclatures, the absence, inaccessibility, or non-transferability of DNA marker information, and the absence of highly contiguous octoploid reference genomes and physically anchored DNA markers (Hardigan et al., 2020, 2021). Those difficulties were eliminated with the assembly and annotation of highly contiguous, haplotype-phased octoploid genomes, the development of octoploid genome-anchored 50K and 850K SNP arrays and medium-density genotyping platforms, the discovery that a significant percentage of short DNA sequences could be unambiguously aligned to octoploid reference genomes, the development of a chromosome nomenclature aligned with the evolutionary history and organization of the four genomes, and the development of a Rosetta Stone for cross-referencing linkage group and chromosome nomenclatures (Supplemental File S3; Tennessen et al. 2014; Hardigan et al. 2020, 2023; Session and Rokhsar 2023).

When DNA sequences for previously genetically mapped *PF*-linked simple sequence repeat (SSR) and amplification-refractory mutation system (ARMS) markers were aligned to the ’UCD Royal Royce’ genome (Sargent et al., 2016; Spigler et al., 2008, 2010; Perrotte et al., 2016; Honjo et al., 2016; Saiga et al., 2022), we discovered that they are located upstream or downstream of the physical position predicted for the *PF* locus in our study (Fig. 6A). Using the available DNA marker sequences, Rosetta Stone, and ’UCD Royal Royce’ genome as a physical reference, we found that at least 10 linkage group nomenclatures have been used to describe the location of DNA markers linked to *PF* on chromosome 4B (Fig. 6A). The linkage group identifiers used in earlier genetic mapping studies are LGIVb-f (Gaston et al., 2013; Perrotte et al., 2016), IV-T-1 (Castro et al., 2015; Honjo et al., 2016), LG-4A (Verma et al., 2017), 4A (Cockerton et al., 2023), LG4-D (Davik et al., 2015), LG4b (Sargent et al., 2016), Fvb4-4 (Edger et al., 2019; Saiga et al., 2022), LGIV (Spigler et al., 2010), LG19 (Spigler et al., 2008), and possibly LG28 (Weebadde et al., 2008). Several of these linkage group nomenclatures, despite variation in acronyms and symbols, were found to have correctly identified chromosome 4, the B genome, or both (Fig. 6A). While the discordance among linkage group nomenclatures can often be reconciled through physical mapping of DNA sequences linked to previously genetically or physically mapped loci, our analysis highlights the critical need for the community-wide adoption of a universal chromosome nomenclature to advance genetic studies in strawberry.

### 4.5 Genome-wide association studies identified a single QTL for runnering

As anticipated we observed LD blocks of *PF*-coincident runner score and count-associated SNPs on the distal end of chromosome 4B in genome-wide association studies of the SDP, F_2_, and full-sib populations (Fig. 5C-H and 6E-G). Those *PF*-associated SNPs identify the *PFRU* QTL previously identified by genetic mapping (Gaston et al., 2013). Significant associations with runner score or count QTL were not observed elsewhere in the genome (Fig. 5). That was unanticipated, especially in the SDP and F_2_ populations. The absence of QTL was less surprising in the full-sib population because the parents were elite individuals with runnering phenotypes that are less extreme than the exotic parents of the F_2_ populations and exotic individuals in the SDP population (Fig. 4; Table 1).

The runnerless phenotypes found in perpetual flowering ecotypes of woodland strawberry (*F. vesca*) are caused by mutations in a single gene (*R*) independent of *S*, a gene that controls the perpetual flowering phenotype in *F. vesca* (Brown and Wareing, 1965). Our initial hypothesis was that the runnerless phenotype of 55C032P001 could have been caused by a single gene mutation independent of the ’Wasatch’ *PF* mutation; however, we found no evidence for that in genome-wide association studies of 17C321P015 × 55C032P001 F_2_ progeny developed by self-pollinating runnerless F_1_ individuals (Fig. 1 and 5; Table 1). Our analyses solely pointed to the *PFRU* QTL (Gaston et al., 2013) and showed that the genetic architecture of runnering is complex in octoploid strawberry (Fig. 8E-G; Table 3).

**TABLE 3.** Statistics^a^ for an A/G SNP marker (AX-184947290) associated with the *PF* locus on chromosome 4B.

| Population<sup>b</sup> | Trait<sup>c</sup> | $\bar{y}_{GG}$ | $\bar{y}_{AG}$ | $\bar{y}_{AA}$ | $\hat{a}$ | $\hat{d}$ | $\vert \hat{d}/\hat{a}\vert $ | PVE (%) |
| --- | --- | --- | --- | --- | --- | --- | --- | --- |
| Diversity Panel | Runner Score | 3.7 | 3.1 | 3.0 | 0.4 | -0.2 | 0.51 | 14.9 |
| Pooled F<sub>2</sub> | Runner Count | 31.5 | 20.8 | 15.0 | 8.3 | -2.4 | 0.29 | 29.1 |
| 21S950 F<sub>2</sub> | Runner Count | - | - | 16.0 | - | - | - | - |
| 21S951 F<sub>2</sub> | Runner Count | 38.6 | 27.9 | 20.5 | 9.1 | -1.7 | 0.19 | 24.6 |
| 21S952 F<sub>2</sub> | Runner Count | 27.7 | 14.2 | 9.9 | 8.9 | -4.6 | 0.51 | 31.1 |
| 21S953 F<sub>2</sub> | Runner Count | 22.1 | 13.7 | 10.8 | 5.6 | -2.8 | 0.49 | 22.6 |
| Full-sib Families | Runner Score | 3.6 | 3.2 | 2.8 | 0.3 | 0.0 | 0.06 | 7.1 |
|  | Runner Count | 50.7 | 37.6 | 28.5 | 11.1 | -2.0 | 0.18 | 15.0 |
|  | Inflorescence Count | 0.0 | 4.5 | 10.6 | -5.3 | -0.8 | 0.14 | 46.7 |

<sup>a</sup> $\bar{y}_{GG}$ , $\bar{y}_{AG}$ , and $\bar{y}_{AA}$ are estimates of SNP marker genotypic means. $\hat{a} = (\bar{y}_{GG} - \bar{y}_{AA})/2$ is the additive effect, $\hat{d} = \bar{y}_{AG} - (\bar{y}_{GG} + \bar{y}_{AA})/2$ is the dominance effect, and $|\hat{d}/\hat{a}|$ is the degree-of-dominance of the AX-184947290 SNP marker locus. PVE is an estimate of the phenotypic variances explained by the SNP marker locus.
<sup>b</sup> Statistics were estimated from clonal replicates of strawberry diversity panel individuals ( $n = 932$ ), unreplicated seed-propagated 17C321P015 × 'Royal Royce' (21S950 and 21S951) and 17C321P015 × 55C032P001 (21S952 and 21S953) F<sub>2</sub> progeny ( $n = 270$ ), and unreplicated seed-propagated full-sib progeny ( $n = 372$ ) developed from nine crosses between elite parents with contrasting runnering phenotypes (Table 1).
<sup>c</sup> The flowering habits of SDP individuals were classified as short-day or day-neutral from the absence or presence of flowering, respectively, under long daylengths (13.5-14.5 hours) in Winters, CA. The runner scores of SDP individuals were recorded over a period of six years in Winters, CA using an ordinal scale, where 1 = runnerless, 2 = weak runnering, 3 = intermediate runnering, 4 = strong runnering, and 5 = extreme runnering (see Fig. 3A). F<sub>2</sub> and full-sib progeny were phenotyped by counting the number of primary runners observed in Davis or Winters, CA, respectively, within one week of the summer solstice (the daylength was 14.5 hours).

The patterns of LD decay for runner score or count-associated SNPs on chromosome 4B differed in the SDP, F_2_, and full-sib populations because of differences in population structure, pedigree inbreeding, recombination, and selection history (Fig. 6E-G; Supplemental Files S6-S8). They provided different pieces to the puzzle for understanding the pleiotropic effects of the *PF* locus on runnering in elite and exotic genetic backgrounds (Table 3). The patterns of LD decay for flowering habit- and runner score-associated SNPs on chromosome 4B were virtually identical in the the highly admixed SDP population (Fig. 4; Supplemental File S6). The lead SNP AX-184947290 was predicted to be closest to the gene encoded by *PF* (Fig. 6E). Using that SNP as a predictor, the *PF* locus was estimated to explain approximately 22.1% of the genetic variance and 14.9% of the phenotypic variance for runner score in the SDP population (Table 3). As observed for flowering habit, when AX-184947290 was used as a fixed effect, the effects of other *PF*-associated SNPs disappeared (Fig. 5D-6E).

The strongest GWAS signals and smoothest patterns of LD decay for runnering were observed in the F_2_ populations (Fig. 5E, 6F, and 8A-D). Fig. 5E and 6F display Manhattan plots for analyses of the F_2_ populations combined. Fig. 8 displays the runner count phenotypic distributions and chromosome 4B Manhattan plots for analyses of individual F_2_ populations. The combined F_2_ population analysis produced sharper and cleaner GWAS signals than those observed in the SDP or full-sib populations because the phenotypes of the parents were extreme and carried unique recombination breakpoints and haplotypic variation was minimum (Table 1; Supplemental Files S7-S8). We observed an LD cliff immediately downstream of AX-184942790 in combined F_2_ and individual 17C321P015 × 55C032P001 F_2_ analyses with comparatively flat LD beyond the LD cliff to the end of the chromosome, in contrast to the choppy pattern of LD decay observed immediately downstream of AX-184942790 in the SDP population (Fig. 6E-F and 8B-D).

We discovered that individuals in the 21S950 (17C321P015 × ’UCD Royal Royce’) F_2_ family were homozygous for ’Wasatch’ alleles from the upper boundary of the ’Wasatch’ introgression (AX-184251339; bp 25,006,055) to AX-184418344 (bp 27,417,624), the 50K array SNP immediately downstream of AX-184947290 (Supplemental File S8). There was an absence of runner-count associated SNPs across the Mb 25.0-30.4 genomic segment in that F_2_ family; hence, we concluded that the gene was likely located upstream of bp 27,417,624 (Fig. 8A). By contrast, significant runner count-associated SNPs were observed across the entire genomic segment (Mb 25.0-30.3) in the other ’17C321P015 × ’UCD Royal Royce’ F_2_ family (21S951). High-density graphical genotypes and haplotypes showed that AX-184947290 and numerous other SNPs upstream of AX-184418344 (bp 27,417,624) segregated in that F_2_ family (Supplemental Files S7-S8). The LD blocks transmitted in those F_2_ individuals were wide, thereby creating a comparatively flat pattern of LD decay across the entire genomic segment in the 21S951 F_2_ family (Fig. 8B).

The runnerless 17C321P015 × 55C032P001 F_1_ individuals that we selected and self-pollinated (19A908P012 and 19A908P053) were discovered to be homozygous upstream of AX-184198749 (bp 26,691,998) and downstream of AX-184947290 (bp 27,294,311) (Supplemental Files S7-S8). That genomic segment segregated in both of those F_2_ families (21S952 and 21S953; Table 1). Fourteen SNPs in the segregating genomic segment were significantly associated with runner count (Fig. 8C-D). Hence, when combined with our association studies in the 17C321P015 × ’UCD Royal Royce’ F_2_ families, we concluded that the gene encoded by *PF* must be located upstream of AX-184418344 (bp 27,417,624) and proximal to AX-184947290 (Supplemental Files S7-S8).

Although the dominant ’Wasatch’ *PF* allele partially suppresses runnering under long days, we found that *PF*_ individuals can have dramatically different runnering phenotypes and that runnering is not always suppressed in *PF*_ individuals, e.g., the runnering phenotype of 21C221P054 (*PFpf*) was extreme (RS = 5.0) and virtually identical to that of the wild parent (Fig. 2-3 and 8-10). At the other extreme, 20C420P006 (*PFpf* was runnerless (RS = 1.0; Fig. 2).

Our analyses of individual F_2_ populations substantiated that phenotypic variation for runnering among short-day (*pfpf*) and day-neutral (*PF*_) individuals cannot be explained by the segregation of *PF* alone (Fig. 2-3 and 8; Table 3). The runner count distributions were strikingly similar among the four F_2_ families; however, phenotypes ranged from runnerless to prolific in three of the four (21S951, 21S952, and 21S953) and weak (10 runners/plant was the minimum) to prolific in the other (21S950) (Fig. 8). GWAS for runner count showed that the 21S950 F_2_ family did not segregate for the *PF* gene, whereas the other three F_2_ families did (Fig. 8; Table 3; (Supplemental Files S7-S8).

The full-sib population combined progeny from nine crosses between elite parents with runnering phenotypes typical of commercially important cultivars (runner scores in the 2.5 to 4.0 range; Table 1; Fig. 10). The individuals in those families were phenotyped for runner and inflorescence count. Association studies of the latter shed light on the pleiotropic effect of the *PF* locus on runner count in elite genetic backgrounds (Fig. 6H). Significant inflorescence count-associated SNPs identify the boundaries of the *PF*-associated LD blocks transmitted by the parents. The pattern of LD decay for flowering habit was comparatively flat from Mb 26.5 to 28.0 among full-sibs because of limited recombination. We observed a distinct LD cliff upstream of *PF* aligned with the stable recombination coldspot found in ’Wasatch’ descendants (Fig. 6H). That pattern clearly shows that LD blocks spanning at least a 0.5 Mb upstream and downstream of the *PF* locus (Mb 27.3) are typically transmitted in segregating populations, partly as a function of limited recombination and partly as a function of shared ancestry (pedigree inbreeding) among elite parents (Table 1) As anticipated, the effect of the *PF* locus was approximately three-fold greater for inflorescence than runner count (Fig. 6G-H; Table 3). Consequently, the LD decay for runner count was comparatively flat and lacked LD cliffs in the full-sib population (Fig. 6G).

We found that runner score was a directionally accurate predictor of runner count, particularly among individuals in the lower tails of both distributions (Fig. 10). The runner counts of seed-propagated full-sib progeny observed near the summer solstice (14.5 photoperiod) were highly correlated with runner score (*r* = 0.86). Notably, the runner count ranges were narrowest among full-sib progeny with runner scores in the runnerless to near-runnerless range (1-2). The runnering phenotypes of at least 18.6% of the full-sib progeny were transgressive (1 ≤ RS ≤ 2). Using the *PF*-associated SNP AX-184947290 as a predictor, of 58 individuals with runner scores = 2, seven were *PFPF*, 29 were *PFpf*, and 22 were *pfpf*; hence, reduced runnering phenotypes were recovered among both short-day and day-neutral individuals.

The statistics reported throughout this paper for runner count were estimated from phenotypes observed when the photoperiod was 14.5 hr, the maximum at our study latitude (Fig. 5-8). To substantiate and model the photoperiod-dependent pleiotropic effect of the *PF* locus on runnering in a typical nursery production (low-elevation clonal propagation) environment (Davis, CA), F_2_ progeny were phenotyped on seven dates beginning 26 March (12.3 hr photoperiod) and ending 6 July 2022 (14.5 hr photoperiod) (Fig. 9). As anticipated, the pleiotropic effect of the *PF* locus on runnering were weakest and non-significant early in the season (26 March to 6 May), increased through the summer solstice (21 June), and was strongest when the photoperiod peaked at 14.5 hr.

### 4.6 SNP haplotypes shed light on the *PF*-associated selective sweep found in ’Wastach’ descendants

The SDP population included 423 day-neutral individuals developed at UC Davis between 1955 and 2021 (’Wasatch’ descendants) and 396 short-day individuals developed at UC Davis between 1935 and 2021 (Fig. 4; Supplemental File S1). With 66 generations of historical recombination (breeding), hybrid offspring of 563 unique pedigrees (fullsib crosses) among 498 parents in the SDP and full-sib populations combined, and highly accurate flowering habit classifications of SDP individuals, we anticipated that GWAS might pinpoint the physical position of the gene encoded by *PF*; however, recombination breakpoints were scarce from the upper boundary of ’Wasatch’ introgression (Mb 26.53) to the lower boundary (Mb 27.28-27.29) of the LD block predicted to harbor *PF* (Fig. 7; Supplemental File S6).

**FIGURE 7.**
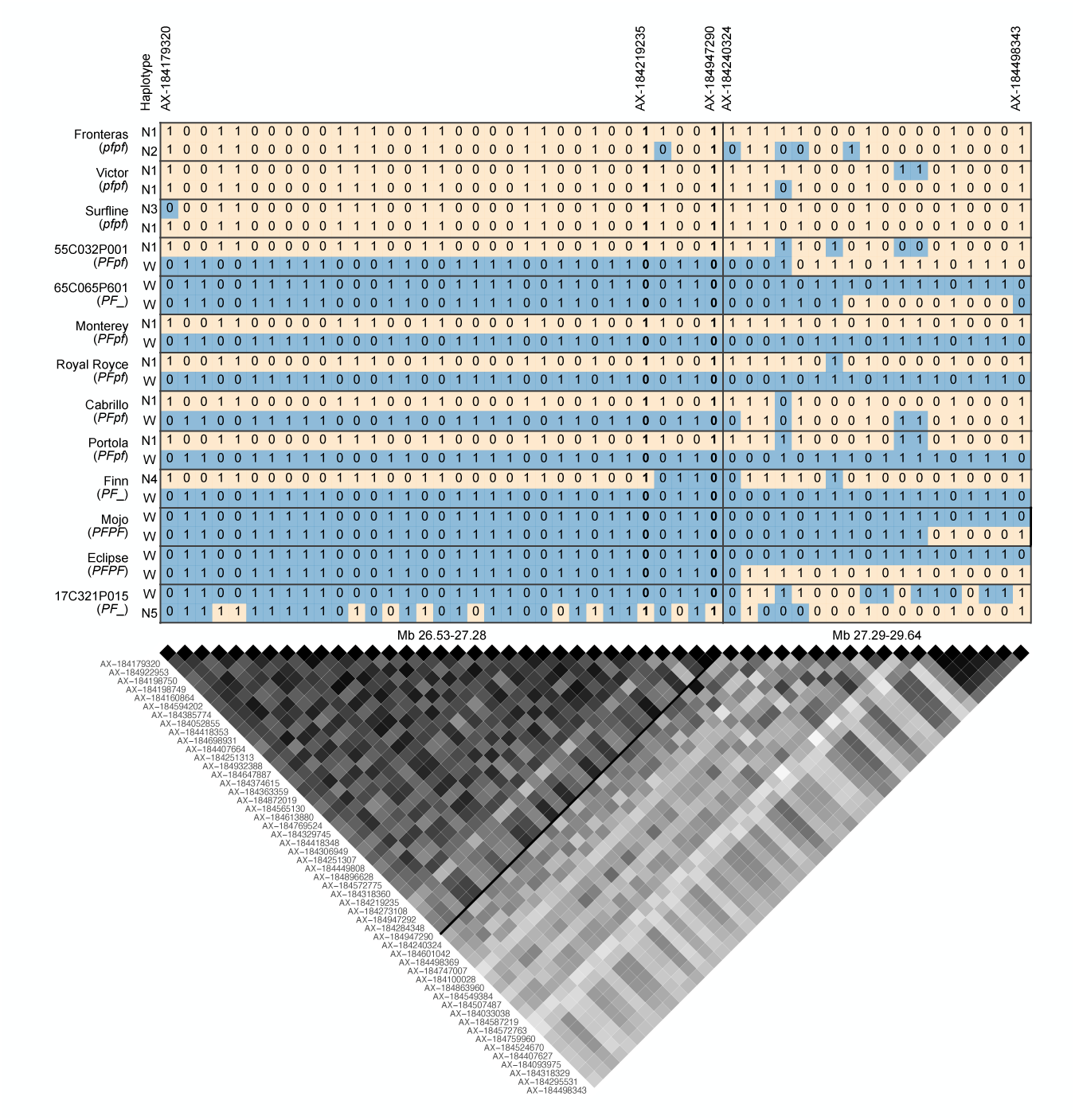
Haplotypes for 51 SNPs flanking the *PF* locus on chromosome 4B (Mb 26.53-29.64) among 808 SDP individuals phenotyped for long day flowering habit. The selected SNPs had minor allele frequencies ≥ 0.3 and exceeded the FDR-corrected significance threshold of 2 for runner score GWAS in the SDP population (−*l og*_10_ *p*-values ≥ 2). The heatmap displays squared correlation coefficient (*R* ^2^) linkage disequilibrium (LD) statistics among the 51 SNPs, where *R* ^2^ = 1 when LD is perfect (black cells), *R* ^2^ = 0 when LD is absent (white cells), and cell darkness increases as *R* ^2^ increases. The 51-SNP LD block was split into a 33-SNP LD block upstream of AX-184240324 and an 18-SNP LD block downstream of AX-184947290 (identified by a solid line between the upper and lower LD blocks). Haplotypes are shown for individuals carrying six out of 400 unique haplotypes in the 33-SNP LD block, including the ’Wasatch’ haplotype (W), a common short-day haplotype (N1), and four others (N2-N5). *PF* genotypes are shown in parentheses below the names of each individual. Axiom 50K array SNP marker names are displayed in the upper panel for a select subset of the 51 SNPs.

**FIGURE 8.**
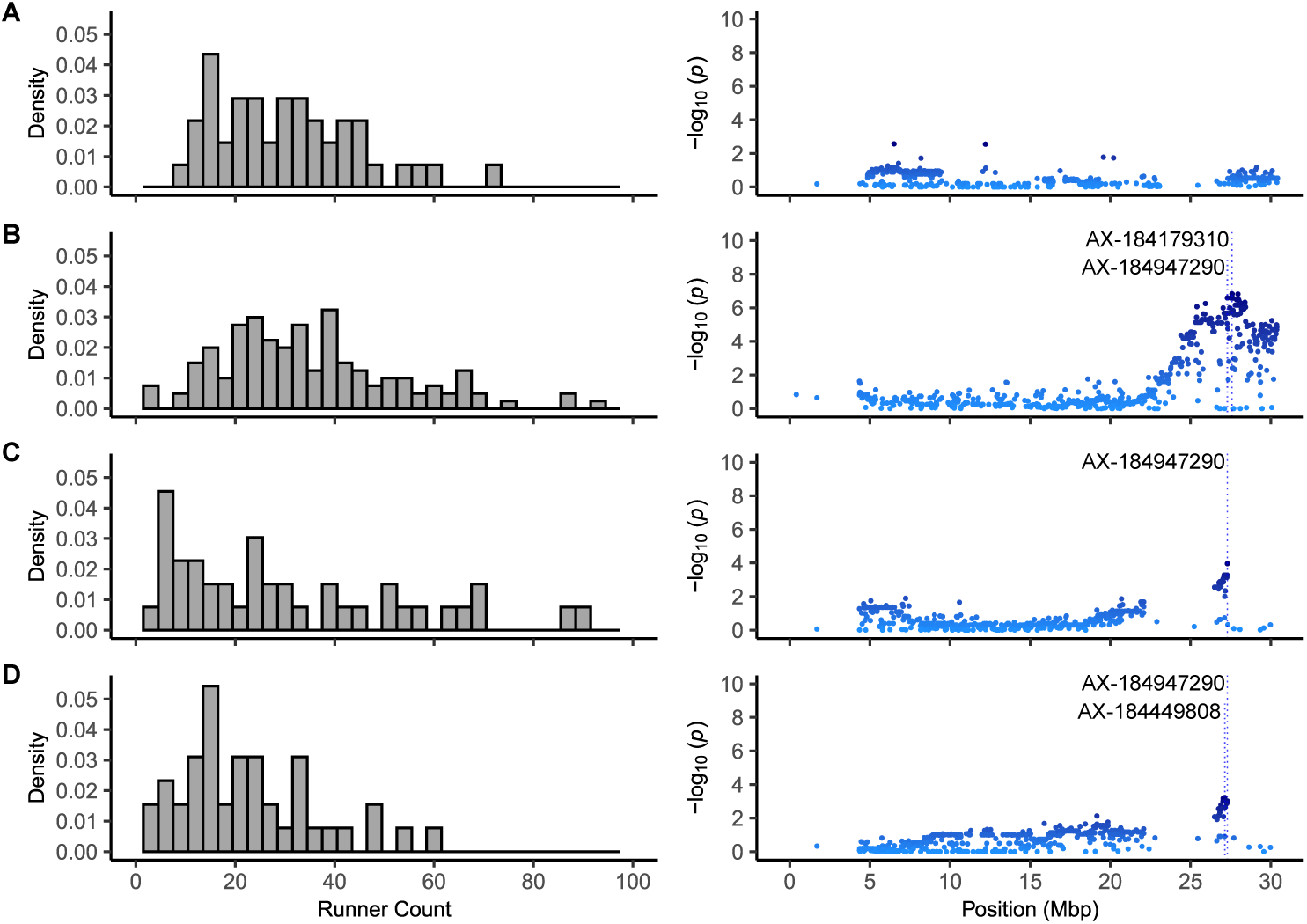
Variation for runner count (left column) and Manhattan plots displaying Axiom 50K array-genotyped SNPs associated with runner count on chromosome 4B (right column) in four F_2_ populations. (A) 17C321P015 × ’UCD Royal Royce’ F_2_ (21S950 originating by self-pollinating the 19A907P022 F_1_). (B) 17C321P015 × ’UCD Royal Royce’ F_2_ (21S951 originated by self-pollinating the 19A907P024 F_1_). (C) 17C321P015 × 55C032P001 F_2_ (21S952 originated by self-pollinating the 19A908P012 F_1_). (D) 17C321P015 × 55C032P001 F_2_ (21S953 originated by self-pollinating the 19A908P053 F_1_). The lead SNPs are labeled.

**FIGURE 9.**
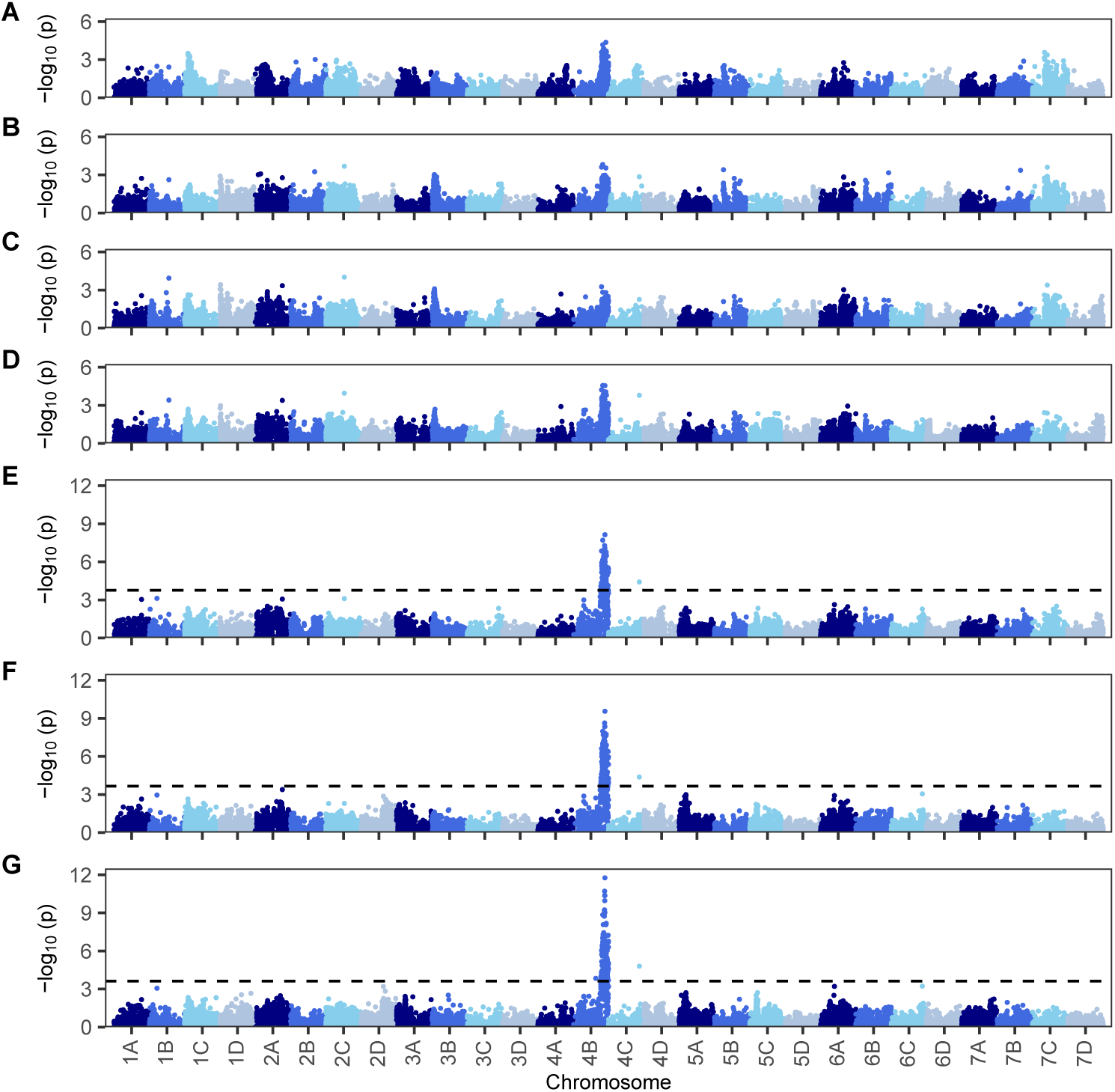
Manhattan plots depicting the photoperiod-dependent effect of the *PF* locus on runner count among 270 F_2_ progeny phenotyped between 26 March and 6 July 2022 in Davis, CA. F_2_ individuals were genotyped with an Axiom 50K SNP array. Manhattan plots are shown for genome-wide association studies of phenotypes observed (A) 26 March (12.3 hr), (B) 9 April (13.0 hr), (C) 22 April (13.3 hr), (D) 6 May (14.1 hr) (E) 20 May (14.3 hr), (F) 3 June (14.1 hr), and (G) 6 July (14.5 hr) among 17C321P015 × ’UCD Royal Royce’ and 17C321P015 × 55C032P001 F_2_ progeny.

**FIGURE 10.**
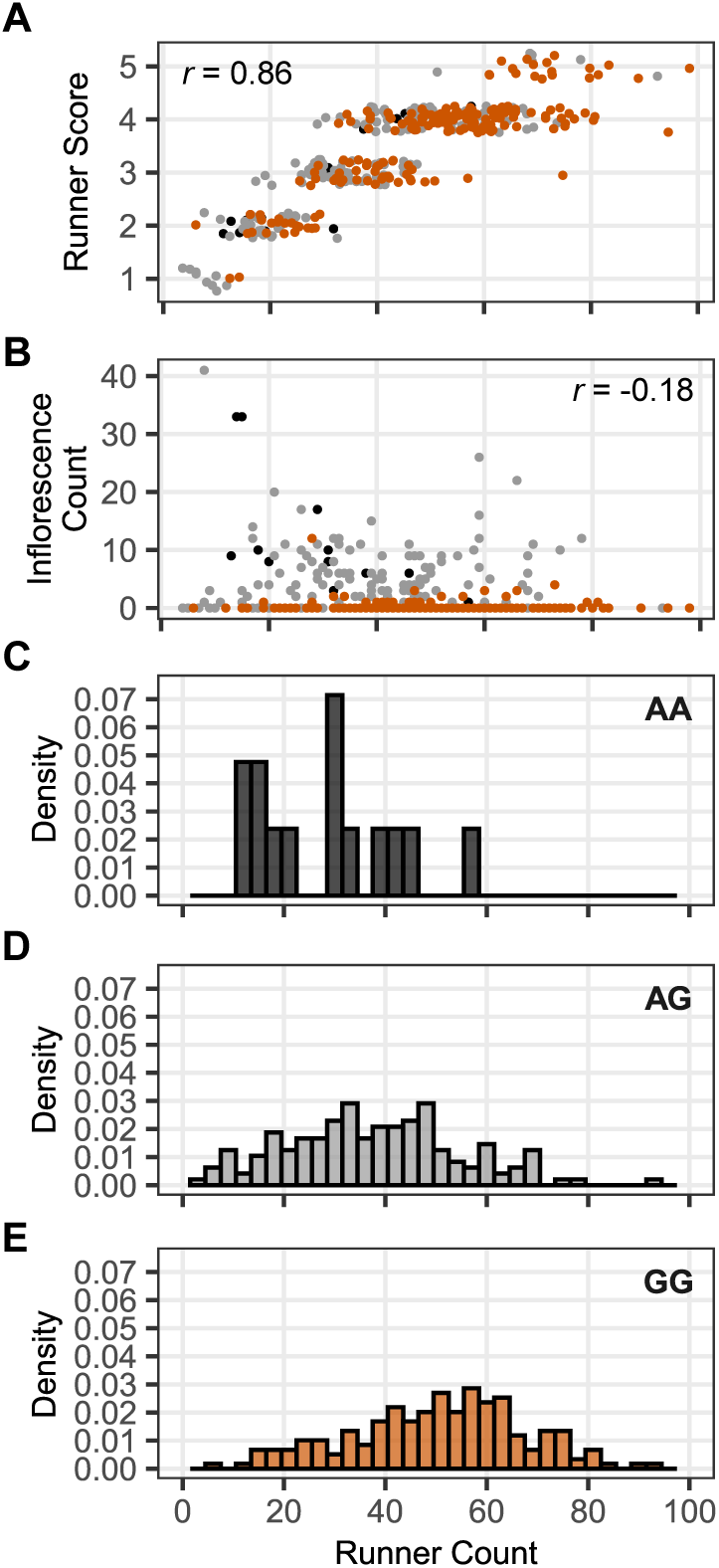
Variation for runner score, runner count, and inflorescence count among 372 full-sib progeny from nine families phenotyped July 1, 2024 in Davis, CA (14.5 hr photoperiod) from seedlings transplanted October 15, 2023. *PF* and the *PF*-associated SNP AX-18497290 segregated in this population. The parents were elite UC short-day and day-neutral individuals (Table 1). (A-E) AX-184947290 SNP genotypes are color coded dark gray (AA), light gray (AG), and orange (GG). (A) The runner count by runner score distribution. (B) The runner count by inflorescence count distribution. (C-E) The runner count distributions among AA, AG, and GG individuals.

To assess the strength and pattern of LD and search for recombination breakpoints, haplotypes of 51 *PF*-associated SNPs were phased among 808 SDP individuals (Fig. 7; Supplemental File S6). The SNPs selected for this analysis had minor allele frequencies (MAFs) greater than 0.3 and GWAS effects for runner score that exceeded a false-discovery-rate (FDR) significance threshold of −*l og*_10_ (p-value) = 2 among SDP population individuals (Fig. 6C-D). We identified 400 unique 51-SNP haplotypes among 808 SDP individuals (1,616 haplotypes; Supplemental File S6). The five most common haplotypes were observed 74 to 234 times each and accounted for 700 of the 1,616 haplotypes (43.3%), whereas 291 haplotypes were only observed once. The 51-SNP ’Wasatch’ haplotype was observed in 87% of the individuals classified as day-neutral (*PF*_).

Strong LD was observed among the 33 phased SNPs upstream of AX-184240324 (Mb 27.29; Fig. 7). LD was markedly weaker among the 18 phased SNPs downstream of AX-184947290 (Mb 27.28). We identified 182 unique 33-SNP haplotypes, including a common ’Wasatch’ haplotype (W) in the upper LD block (Fig. 7). The boundary between the upper and lower LD blocks was supported by the absence of *PF* segregation among 17C321P015 × 55C032P001 F_2_ progeny and among full-sib progeny developed from crosses between ’UC Eclipse’ and short-day cultivars. ’UC Eclipse’ carries a recombination breakpoint immediately downstream of AX-184240324; hence, we concluded that *PF* is found upstream of the upper boundary of the lower LD block. ’Cabrillo’ and ’UCD Finn’ share that recombination breakpoint (Fig. 7). When combined with insights gained from unique recombination breakpoints among F_2_ progeny, SNP haplotypes of SDP individuals, and the presence or absence of *PF* segregation among F_2_ progeny and ’UC Eclipse’ × short-day full-sib progeny, we are confident that the *PF* gene is found between AX-184021395 (Mb 26.40) and AX-184240324 (Mb 27.28) on chromosome 4B (Fig. 6-8; Supplemental Files S6-S8).

Our search of annotations in ’UCD Royal Royce’ genome assemblies (both haplotypes) identified 193 genes spanning bp 26,401,121 to 27,418,663 on chromosome 4B (Supplemental File S9). That search did not uncover any genes known or predicted to regulate flowering in angiosperms (Hempel et al., 1997). We did, however, discover a homolog of *REPRESSOR OF GA1* on chromosome 4B (*RGA1-4B*) downstream of *PF*. *RGA1-4B* gene identifiers in the haplotype-phased ’UCD Royal Royce’ genome assemblies are Fxa4Bg203107 (haplotype 2; Mb 28.83-28.94) and Fxa4Bg103085 (haplotype 1; Mb 29.15-29.25). Homoeologs were also identified in the other three genomes (Supplemental Fig. S1). Studies in *F. vesca* identified a *REPRESSOR OF GA1* (*FveRGA1*) gene on chromosome 4 as a causal determinant of the runnerless phenotype of ’Yellow Wonder’ (Tenreira et al., 2017; Caruana et al., 2018). ’Hawaii 4’ develops runners under long days, whereas ’Yellow Wonder’ does not.

The *PF*-associated SNPs identified in the present study and *PF*-associated SSR and ARMS markers markers identified in previous studies are imperfect predictors of *PF* genotypes because of recombination history, haplotype complexity, and admixture (Perrotte et al., 2016; Salinas et al., 2017; Honjo et al., 2016; Saiga et al., 2022). Several array-genotyped SNPs in the upper LD block predicted the flowering habit of SDP individuals with similar accuracy. The A allele for the AX-184947290 SNP (found in the ’Wasatch’ haplotype) was observed in 332 out of 379 SDP individuals classified as day-neutral (87.6%). The AX-184947290 GG homozygote was observed in 345 out of 416 SDP individuals classified as short-day (82.9%). The prediction accuracy of AX-184947290 was marginally better when non-UC individuals were trimmed from the analysis (88.5% for DN and 86.7% for SD individuals). That trend was repeated for several other individual SNPs and for 3- and 5-SNP haplotypes in the upper LD block, e.g., the 33-SNP ’Wasatch’ haplotype (W) was observed in 84.6% of the the SDP individuals classified as day-neutral (*PF*_) in the SDP population, including 55C031P001, 65C065P601, and UC day-neutral and summer-plant cultivars (Supplemental File S6). The obvious solution to the imperfect prediction problem is to identify the causal gene and mutation underlying *PF*. Despite extensive research, and the >900 individuals and >60 generations presented here, this causal variant still remains elusive.

Tenreira et al. (2017) showed that the runnerless phenotype of ’Yellow Wonder’ was caused by “a deletion mutation in a gene coding for a GA20-oxidase enzyme needed for GA biosynthesis”. Caruana et al. (2018) induced a recessive mutation (*suppressor of runnerless-1*; *srl-1*) in Yellow Wonder that eliminated the runnerless phenotype. The *srl-1* phenotype was subsequently found to be caused by a point mutation (G → A) that created a stop codon in *FveRGA1*, *FveRGA1*, which encodes a *DELLA* protein. They showed that the mutation affects the relative expression of *FveRGA1* in leaves and concluded “that *FveRGA1* likely represses GA signaling in multiple tissues and developmental stages” and that *FveRGA1* plays an important role in “regulating axillary meristem identity in strawberry” (Caruana et al., 2018). We originally hypothesized that runnerless phenotype of 55C032P001 and quantitative genetic variation for runnering in octoploid strawberry might be caused by allelic variation of an *RGA1* homoeolog. While the association between *RGA1-4B* and *PF* was intriguing, the LD was weak and the effects of *RGA-4B*-associated SNPs on runner score or count were weak (Supplemental Fig. S1).

### 4.7 The pleiotropic effect of the *PERPETUAL FLOWERING* locus on runnering across a broad cross-section of octoploid genetic resources

Our data support the hypothesis that the *PFRU* QTL for runnering previously identified by Gaston et al. (2013) was likely caused by the pleitropic effect of a single gene (*PF*). Using the lead SNP identified by GWAS (AX-184947290), we estimated that the *PF* locus explained 7.1 to 31.1% of the phenotypic variation for runner score or count (Table 3). The ’Wasatch’ allele has a dominant (Mendelian) effect on long day flowering habit; however, the pleiotropic effects of the *PF* allele on runnering are quantitative and incompletely dominant: |*d*^^^/*a*^| ranged from 0.18 in the elite × elite full-sib population to 0.51 in the SDP population (Table 3).

The *PF* locus is well known to affect the timing of the photoperiod-dependent inhibition of flowering and discrete transition from sexual to asexual reproduction in short-day (*pfpf*) plants (Ahmadi et al., 1990; Gaston et al., 2013; Sønsteby and Heide, 2006). While neither are observed in day-neutral (*PF*_) plants (Gaston et al., 2013; Sønsteby and Heide, 2006), both short-day and day-neutral plants runner under a wide range of conditions, including short and long days, and when producing fruit, which is why runner trimming is commonly practiced and necessary to promote flowering and maximize yield in fruit production (Fig. 1-3). We observed the complete range of phenotypes, from runnerless (RS = 1.0) to extreme runnering (RS = 5.0), among the diverse sample of SD and DN individuals assembled for the SDP and parents of F_2_ families segregating for *PF* alleles and runnering phenotypes (Table 1; Fig. 3; Supplemental File S1).

The runnering phenotypes of most of the wild species ecotypes in our study were aggressive (RS > 4.0), as exemplified by Frederik 9, Hinesburg, Harris Spring, and Yaquina A in Fig. 1 (Supplemental File S1). The median runner score was *ỹ* = 4.8 among *F. chiloensis* and *ỹ* = 4.8 among *F. virginiana* ecotypes. The runner score means for 33 of the 38 octoploid ecotypes phenotyped in our study were in the 4.0 to 5.0 range, and only one (*ȳ* = 3.0; Scott’s Creek) was less than the SDP population mean (*ȳ* = 3.38).

The runner score distribution was negatively-skewed among short-day and approximately normal among day-neutral individuals in the SDP population (Fig. 3). The runner score mean was significantly greater for short-day (*pfpf*) individuals (*ȳ_SD_* = 3.68) than day-neutral (*PF*_) individuals (*ȳ_DN_* = 3.07; *p* ≤ 0.0001). Similarly, the runner score median was greater for SD individuals (*ỹ_SD_* = 3.95) than DN individuals (*ỹ_DN_* = 2.97) (Fig. 3; Supplemental File S1). These differences broadly reflect the pleiotropic effect of the *PF* locus on runnering over an entire growing season in our low-elevation nursery production (clonal propagation) environment (Fig. 1-2).

As shown for a broad cross-section of octoploid genetic resources, DN (*PF_*) individuals tend to produce fewer runners than SD (*pfpf*) individuals in nursery production (Fig. 3; Supplemental File S1). Of 478 SD individuals in the SDP population, 97.5% had runner scores in the 2.5-5.0 range. The 12 SD individuals in the weak runnering range (1.0-2.5) include the cultivars ’Elsanta’ (RS = 1.2), ’Howard 17’ (RS = 2.0), ’Senga Sengana’ (RS = 2.0), and ’Fairfax’ (RS = 2.0). The other eight are UC individuals developed in 2018 that were purposefully selected for reduced runnering (Supplemental File S1).

We observed a wide range of runnering phenotypes among DN individuals known to be homozygous or heterozygous for the dominant ’Wasatch’ *PF* allele, e.g., 20C420P006 (*PFpf*; RS = 1.0), ’UCD Moxie’ (*PFpf*; RS = 2.0), ’UC Eclipse’ (*PFPF*; RS = 3.4), and ’Monterey’ (*PFpf*; RS = 4.0) (Fig. 2). Of 444 individuals classified as day-neutral in the SDP population, 80.0% had runners scores in the 2.5 to 5.0 range (Fig. 3; Supplemental File S1). The percentage of individuals in the weak runnering range (1.0-2.5) was greater for DN (20.0%) than SD (2.5%) individuals. Weak runnering DN cultivars were uncommon (the runnering phenotypes of 30 of the 40 DN cultivars screened in our study were intermediate to strong).

DN cultivars carrying the ’Wasatch’ *PF* allele have occasionally been described as poor runner producers, but cannot be broadly classified as poor runnering, e.g.,Hossain et al. (2019) described ’Albion’ (*PFpf*) as a “poor runner producer” and depicted a runnerless ’Albion’ plant. Our data do not support that characterization. ’Albion’ produced an above average number of runners (RS = 3.3), runners in fruit and bare-root plant production environments, and has been clonally propagated on a large-scale for commercial production since 1997 (Fig. 2; Supplemental File S1). Our data show that the runnering phenotypes of day-neutral (*PF*_) individuals temporally vary and span the entire range, from runnerless to extreme (Fig. 3, 8-10).

### 4.8 Accuracy of genomic selection for runnering

The *PF* locus was found to be beneficial but insufficient for predicting runnering phenotypes in strawberry populations segregating for dominant and recessive *PF* alleles (Table 3; Fig. 1-2). Moreover, the pleiotropic effect of the *PF* locus on runnering is inconsequential in cultivar development because the selected individuals are destined to either be short-day (*pfpf*) or day-neutral (*PF_*). To state this another way, when selecting for runnerless or reduced runnering phenotypes within short-day or day-neutral populations *per se*, the outcome of selection hinges on genetic variation caused by loci other than *PF* (Table 3). Those other loci underlie the missing heritability in our study.

With as much as 78% of the heritability for runnering missing (unexplained by segregation of the *PF* locus), we explored the merit of applying genomic selection for runnering by testing different combinations of training and validation populations with and without correcting for the fixed effect of the *PF* locus (Table 4). We specifically sought to identify how and where genomic selection for runnering might be cost effectively applied in a strawberry breeding pipeline to identify runnerless individuals as selection candidates for seed-propagated cultivars and near-runnerless individuals as selection candidates for clone-propagated cultivars. The strategy for the latter is to identify individuals near the lower limit of commercially viable clonal propagation, specifically those with runner scores in the 2.0-2.5 range (Fig. 3 and 10).

**TABLE 4.** Genomic predictive ability (*r_a_*_^,*ȳ*_) and prediction accuracy (*r_a_*_^,*ȳ*_ *h*) for runner score or count within and between strawberry populations estimated by cross-validation^a^ with and without correcting for the effect of the *PF* locus^b^.

| Population<sup>c</sup> |  | PF Uncorrected |  | PF Corrected |  |
| --- | --- | --- | --- | --- | --- |
| Training | Validation | $r_{\hat{a},y}$ | $r_{\hat{a},y}/h$ | $r_{\hat{a},y}$ | $r_{\hat{a},y}/h$ |
| SDP | SDP | 0.59 | 0.76 | 0.60 | 0.77 |
| SDP-UC | SDP-UC | 0.47 | 0.62 | 0.50 | 0.66 |
| FS | FS | 0.55 | 0.75 | 0.58 | 0.79 |
| SDP | FS | 0.33 | 0.45 | 0.43 | 0.60 |
| SDP-UC | FS | 0.25 | 0.34 | 0.39 | 0.53 |

<sup>a</sup> Cross-validation was done by randomly drawing 80% of the individuals within a training population to predict genomic-estimated breeding values ( $\hat{a}$ ) of validation population individuals. The prediction accuracy was estimated by $r_{\hat{a},y}/h$ , where $r_{\hat{a},y}$ is the simple correlation between the observed phenotype ( $y$ ) and $\hat{a}$ , $h = \sqrt{h^2}$ , $h^2$ is the genomic-estimated narrow-sense heritability, $y$ was the either the runner count for unreplicated seed-propagated full-sib (FS) individuals or the estimated marginal mean for runner score for clonally propagated strawberry diversity panel (SDP) individuals over years. Cross-validation statistics were estimated from 1,000 randomly drawn samples of training population individuals with replacement. When predicting within a population (e.g., within the SDP population), $\hat{a}$ was estimated for individuals that were not sampled in each iteration (20% of the individuals within the population). When predicting between populations, $\hat{a}$ was estimated for every individual in the validation population.
<sup>b</sup> The *PF*-associated SNP AX-184947290 was used as a fixed effect to correct for the effect of the *PF* locus on runner score or count.
<sup>c</sup> The populations are the strawberry diversity panel (SDP; $n = 932$ ), a population-specific subset of cultivars and other elite strawberry diversity panel individuals developed at the University of California, Davis (SDP-UC; $n = 672$ ), 17C321P015 × 'Royal Royce' and 17C321P015 × 55C032P001 F<sub>2</sub> families ( $n = 270$ ), and nine full-sib families ( $n = 372$ ) developed from crosses between elite UC parents.

Because our training populations (SDP and full-sib, FS) segregated for dominant and recessive *PF* alleles, we used a *PF*-associated SNP marker (AX-184947290) to correct for the fixed effect of the *PF* locus (Tables 1 and 4). The inclusion of AX-184947290 as a fixed effect increased the accuracy of genomic prediction across the training and validation population combinations tested (Table 4). The accuracy increase gained by *PF*-correction was greater for the between population combinations (training → validation = SDP → FS and SDP-UC → FS) than the within population combinations tested in our study, e.g., SDP → SDP (Table 4). As anticipated, directly correcting for the fixed effect of the *PF* locus is necessary for maximizing genomic prediction accuracy when selecting for runnering in populations segregating for dominant and recessive *PF* alleles (Table 4).

The clonal genetic resources associated with strawberry breeding programs are typically comprised of permanent to semi-permanent individuals, both exotic and elite, and a continually evolving sample of selected individuals spanning several generations of within full-sib family selection. The 932 individuals selected for the SDP originated from 554 unique pedigrees (full-sib families) among 489 parents (Supplemental File S1). The phenotypic data underlying breeding program-associated clonal genetic resources are continually updated, thereby creating ever-increasing samples of training population data to inform parent and genomic selection (Supplemental File S1). We used the SDP to assess whether runner score observations accumulated year-over-year among genetically diverse individuals could be used to accurately estimate breeding values for runner count among seed-propagated elite × elite full-sib progeny (selection candidates before clonal propagation). We discovered that breeding values for runner score were predicted with outstanding accuracy within the SDP population *per se* (the *PF*-corrected estimate was *r_a_*_^,*ȳ*_ /*h* = 0.77). The prediction accuracy was lower when the SDP was used as the training population for predicting breeding values among elite × elite full-sib progeny (SDP → FS; Table 4). It must be noted here that (i) the SDP plants were runner-propagated bare-root clones while the full-sibs were seedlings, and (ii) the SDP plants were scored 1-5, while the full-sibs were phenotyped with a runner count. Prediction accuracy was reasonable despite these physiological and methodological differences.

To explore the SDP approach further, we tested a training population comprised of 776 SDP individuals with UC pedigrees starting from 1969 onward (SDP-UC), the year of origin of the first ’Wasatch’-derived day-neutral cultivars, specifically ’Hecker’, ’Aptos’, and ’Brighton’ (Bringhurst and Voth, 1980). The SDP-UC subset was created by eliminating 156 pre-1969 UC, non-UC, and other exotic individuals from the SDP (Supplemental File S1). Because accuracy depends on the closeness of genetic relationships and patterns of linkage disequilibrium between training and validation populations (Habier et al., 2007; Lorenz and Smith, 2015; Hickey et al., 2014), our hypothesis was that accuracy could be increased by using a UC-specific subset of SDP training population individuals to predict the breeding values of full-sib progeny developed with elite UC parents. We discovered that runner score breeding values were predicted with acceptable accuracy within the SDP-UC training population *per se* (SDP-UC → SDP-UC); to our surprise however, genomic prediction accuracy was lower within the SDP-UC than the SDP training population, e.g., *r_a_*_^,*ȳ*_ /*h* decreased from 0.77 in the SDP → SDP cross-validation to 0.66 in the SDP-UC → SDP-UC cross-validation (Table 4). The genomic prediction accuracy achieved with the SDP training population was superior to that of the SDP-UC training population in both of the between population scenarios tested: SDP → FS and SDP-UC → FS (Table 4).

We attributed the decreased accuracy of the SDP-UC training population partly to the reduction of individuals from the upper and lower tails of the runner score distribution and associated decreases in allelic diversity among loci underlying extreme phenotypes eliminated by artificial selection (Fig. 10). While adding genetically distant individuals to a training population often decreases genomic prediction accuracy, we suspect that the effect of eliminating genetically distant individuals from the training population was offset by the loss of allelic variation needed for accurate genomic prediction (Habier et al., 2007; Lorenz and Smith, 2015; Hickey et al., 2014). Selection against runnering phenotypes at both extremes (RS < 2.0 and RS > 4.0) has narrowed genetic variation for runnering among modern cultivars, including those in the SDP-UC subset. Our analysis suggests that augmenting the training population with extreme phenotypes can increase prediction accuracy, similar to what has been proposed in similar breeding scenarios where selection has significantly changed allele frequencies and eliminated unfavorable alleles (Brandariz and Bernardo, 2018; Knapp et al., 2024).

The prediction accuracy for runner count within the full-sib training population (FS → FS) was nearly identical to that observed within the SDP training population (Table 4). This suggested that phenotyping a nominal number of seed-propagated (unreplicated) full-sib individuals for runner count should facilitate the accurate prediction of breeding values of unphenotyped full-sib individuals, particularly with continual updating of the training population (Jannink et al., 2010; Hickey et al., 2014; Lorenz and Nice, 2017). The approach of counting runners among a subset of individuals within full-sib families, although effective for building a training population and applying genomic selection, is time consuming, cost prohibitive, and unnecessary for identifying individuals with runnerless to near-runnerless phenotypes, primarily because rapid visual phenotyping and phenotypic selection among seed-propagated full-sib individuals was found to be highly effective (Fig. 1-2).

### 4.9 Limitations and Conclusions

Our studies were undertaken to assess the feasibility of identifying runnerless and reduced runnering selection candidates in segregating populations. Although the genetic mechanisms underlying variation for runnering appear to be complex (a quantitative black box), the broad-sense heritability, stability, and reproducibility of runnering phenotypes were exceptional and showed that the natural genetic variation needed to develop runnerless or near-runnerless cultivars is prevalent and heritable in octoploid strawberry. We recovered runnerless and near-runnerless phenotypes at moderately high frequencies and found them to be stable and reproducible despite the complex genetic architecture of runnering.

Deeper studies than those presented here are needed to understand how runnerless and near-runnerless phenotypes affect yield, canopy architecture, and plant development. We hypothesized that runnering and fruit yield are negatively genetically correlated and posited that reduced runnering phenotypes should increase fruit yield, in addition to greatly decreasing runner trimming costs for fruit growers. The latter has been substantiated by large-scale commercial testing of ’UCD Moxie’, a prototypical reduced runnering, day-neutral cultivar that produces high yields of fruit and requires substantially less runner trimming than standard day-neutral cultivars (Fig. 2). The pleiotropic effect of runnering on fruit yield, however, has not yet been commercially documented because runners are mechanically trimmed in production fields, thereby masking the negative pleiotropic effect of runner and daughter plant growth on flowering and fruit yield.

Our hypothesis that the runnerless phenotype of 55C032P001 was caused by a single gene mutation was not accepted. While the pleiotropic effect of *PF* on runnering was predictable and anticipated (Gaston et al., 2013), the absence of large-effect loci for runnering was not. The identification of single gene mutations with large effects on runnering in future studies seems plausible, particularly since *RGA1* and other genes affecting flowering and runnering have been identified and others are bound to emerge (Koskela et al., 2016; Caruana et al., 2018; Hytönen and Kurokura, 2020). We identified a homolog of one of the more promising candidate genes (*RGA1*) in the fringe of the *PF* LD block (*RGA1-4B*; Fxa4Bg103085; bp 28,885,347-28,887,164). While intriguing, SNPs in LD with *RGA1-4B* were 1.59 Mb downstream of the most likely location of *PF* (27,294,311) and only very weakly associated with runner count. Whether natural *RGA1-4B* allelic variants affect runnering remains unknown. That gene is clearly an interesting target for genetic engineering or gene editing (Caruana et al., 2018).

Strawberry production could evolve, at least partly, towards seed-propagated cultivars where runners are unnecessary and diminish productivity by increasing labor (runner trimming) costs and diverting energy and nutrients away from sexual to asexual reproduction. This necessitates a shift from the *status quo* of clonal propagation of hybrid individuals between outbred (partially inbred) parents to seed propagation of single-cross hybrids between inbred parents, the creation and exploitation of heterotic groups, and the development of a hybrid seed production system (Feldmann et al., 2024a). Seed propagation of hybrid individuals through apomixis could be even more game-changing (Conner and Ozias-Akins, 2017; Underwood and Mercier, 2022; Mahlandt et al., 2023), eliminate the need for implementing resource-intensive inbred-hybrid breeding schemes, and facilitate less resource-intensive within family selection schemes that have produced substantial genetic gains and can directly exploit elite genetics without inbred line development (Feldmann et al., 2024a,b).

## AUTHORS CONTRIBUTIONS

**Hillel Brukental:** Data curation; formal analysis; funding acquisition; investigation; methodology; validation; visualization; writing – original draft; and writing – review & editing. **Marta L. Bjornson:** Conceptualization; data curation; formal analysis; investigation; methodology; supervision; validation; visualization; and writing – review & editing. **Dominique D.A. Pincot:** Data curation; formal analysis; methodology; and writing – review & editing. **Michael A. Hardigan:** Data curation; formal analysis; investigation. **Sadikshya Sharma:** Data curation; investigation. **Nicolas P. Jimenez:** Investigation. **Randi A. Famula:** Data curation; formal analysis; funding acquisition; investigation; methodology; project administration; resources; software; supervision; validation; visualization; writing – original draft; and writing – review & editing. **Cindy M. Lopez Ramirez:** data curation; investigation. **Glenn S. Cole:** conceptualization; data curation; investigation; methodology; project administration; and writing – review & editing. **Mitchell J. Feldmann:** Cconceptualization; data curation; formal analysis; funding acquisition; investigation; methodology; project administration; supervision; validation; visualization; writing – original draft; and writing – review & editing. **Steven J. Knapp:** Conceptualization; formal analysis; funding acquisition; investigation; methodology; project administration; Resources; supervision; validation; visualization; writing – original draft; and writing – review & editing.

## ACKNOWLEDGEMENTS

This research was supported by a Vaadia-BARD Postdoctoral Fellowship awarded to HB by the United States-Israel Binational Agricultural Research and Development Fund (Award #FI-637-2023) and by grants awarded to SJK and MJF from the United Stated Department of Agriculture (http://dx.doi.org/10.13039/100000199) National Institute of Food and Agriculture (NIFA) Specialty Crops Research Initiative (#2017-51181B6833 and #2022-51181-38328-0) and California Strawberry Commission (http://dx.doi.org/10.13039/100006760). We greatly appreciate the outstanding support of the field superintendents and staff at the UC Davis Wolfskill Experiment Orchard (Winters, CA), and UC Davis Vegetable Crops (Davis, CA).

## Supporting Information

The phenotypic and genotypic data generated in this study have been deposited at Zenodo and are freely available (https://doi.org/10.5281/zenodo.13937029).

**Supplemental File S1.** Origin years, species, pedigrees, flowering habit classifications, runner score phenotypic means, and AX-184947290 SNP genotypes for *n* = 932 octoploid strawberry diversity panel (SDP) individuals. The SDP included 15 *F. chiloensis*, 24 *F. virginiana*, and 893 *F.* × *ananassa* clonally propagated individuals. SDP individuals were classified as short-day (SD = 0) or day-neutral (DN = 1). The runner scores of SDP individuals were recorded on clonally propagated plants in Winters, CA using an ordinal scale, where 1 = runnerless, 2 = weak runnering, 3 = intermediate runnering, 4 = strong runnering, and 5 = extreme runnering. The runner score estimated marginal means (EMMs) of SDP individuals were estimated from *r* = 1.92 observations, where *r* is the harmonic mean number of observations per individual (the data were unbalanced and some individuals were observed only once). Tabs 2 and 3 present the unique crosses (554) and unique parents (489) of the 932 SDP individuals.

**Supplemental File S2.** The physical positions of 50K and 850K Axiom array SNPs corroborated by genetic mapping. SNPs were physically anchored to the ’Camarosa’ (FaCA1; Edger et al. (2019); https://phytozome-next.jgi.doe. gov/info/Fxananassa_v1_0_a1) and ‘UCD Royal Royce’ (FaRR1; https://phytozome-next.jgi.doe.gov/info/ FxananassaRoyalRoyce_v1_0) genomes *in silico*. Chromosomes were numbered using the nomenclature of Hardigan et al. (2020) and cross-referenced to the nomenclature of Edger et al. (2019). This database includes the physical positions of SNPs identified by BLAST in the ’Camarosa’ and ’Royal Royce’ genomes, DNA sequences of the SNP probes, chromosome assignments and physical positions of SNPs corroborated by genetic mapping, and associated information.

**Supplemental File S3.** A Rosetta stone for cross-referencing linkage group and chromosome nomenclatures in octoploid strawberry. This database includes several previously published chromosome nomenclatures (Tennessen et al., 2014; Edger et al., 2019; Hardigan et al., 2020) and the chromosome nomenclature proposed by Session and Rokhsar (2023).

**Supplemental File S4.** Pedigrees, runner count phenotypes, and AX-184947290 SNP genotypes for *n* = 87 seed-propagated 17C321P015 × 55C032P001 F_2_ and *n* = 183 seed-propagated 17C321P015 × ’UCD Royal Royce’ F_2_ progeny observed 26 March, 9 April, 22 April, 6 May, 20 May, 3 June, and 6 July 2023 in Davis, CA (the respective photoperiods on those dates were 12.3, 13.0, 13.3, 14.1, 14.3, 14.1, and 14.5 hr). Four F_2_ families (21S950, 21S951, 21S952, and 21S953) were developed by self-pollinating runnerless to near-runnerless F_1_ individuals (19A907P022, 19A907P024, 19A908P012, and 19A908P053) originating in 17C321P015 × 55C032P001 and 17C321P015 × ’UCD Royal Royce’ full-sib families (Table 1).

**Supplemental File S5.** Pedigrees, runner count, runner score, and inflorescence count phenotypes, and AX-184947290 SNP genotypes for 372 seed-propagated full-sib progeny observed 1 May, 1 June, and 1 July 2024 in Winters, CA (the respective photoperiods on those dates were 13.5, 14.4, and 14.5 hr). The population included nine full-sib families developed from crosses among 13 elite UC parents (Table 1).

**Supplemental File S6.** Haplotypes for 51 phased SNPs among 808 SDP individuals. This database includes 1,606 0,1-coded haplotypes among 808 SDP individuals classified as either short-day (SD = 0) or day-neutral (DN = 1).

**Supplemental File S7.** Graphical genotypes for 50K Axiom array-genotyped SNPs segregating between Mb 25.0 and 30.4 on chromosome 4B in 17C321P015 × 55C032P001 and 17C321P015 × ’UCD Royal Royce’ F_2_ families. This database includes runnering phenotypes, SNP genotypes, and the ordered physical positions for 140 SNP loci among 279 individuals within four F_2_ families (21S950, 21S951, 21S952, and 21S953). Genotypes are colored by their PFRU associations: peach = *pf/pf*, pale green = *PF/pf*, darker green = *PF/PF*.

**Supplemental File S8.** Genotypes and haplotypes for 15 SNPs associated with the *PF* locus on chromosome 4B among selected samples of three individuals each from an 17C321P015 × ’UCD Royal Royce’ F_2_ family (21S950) and two 17C321P015 × 55C032P001 F_2_ families (21S952 and 21S953). The SNPs spanned bp 26,389,744 (AX-184068183) to bp 27,597,497 (AX-184160850). Genotypes are colored by their PFRU associations: peach = *pf/pf*, pale green = *PF/pf*, darker green = *PF/PF*.

**Supplemental File S9.** The predicted functions of 173 annotated genes spanning bp 27,597,497 to 27,418,663 on chromosome 4B in the ’UCD Royal Royce’ reference genome (FaRR1; https://phytozome-next.jgi.doe.gov/ info/FxananassaRoyalRoyce_v1_0).

**Supplemental Figure S1.** Synteny of *REPRESSOR OF GA1* (*RGA1*) homoeologs on chromosome 4 in the A, B, C, and D genomes of octoploid strawberry (*F.* × *ananassa*) and A genome of woodland strawberry (*F. vesca*). The ortholog identified in *F. vesca* (*FveRGA1*) by Caruana et al. (2018) is shown below *F.* × *ananassa* chromosome 4 haplotypes. Homoelogs found in the octoploid are identified by the prefix Fxa. Homology was calculated using GENESPACE Lovell et al. (2022), and the synteny plot was generated using JCVI Tang et al. (2024). The synteny plot was developed using genes annotated in the haplotype-phased assemblies of the ’UCD Royal Royce’ genome, a day-neutral *F.* × *ananassa* cultivar (https://phytozome-next.jgi.doe.gov/info/FxananassaRoyalRoyce_v1_0.)

## Abbreviations

ARMS: amplification-refractory mutation system
DN: day-neutral
EMM: estimated marginal mean
FH: flowering habit
FS: Full sibs
GWAS: genome-wide association study
GEBV: genomic-estimated breeding value
LMM: linear mixed model
LD: linkage disequilibrium
MAS: marker-assisted selection
PF: perpetual flowering
QTL: quantitative trait locus
REML: restricted maximum likelihood
RC: runner count
RS: runner score
SD: short-day
SNP: single nucleotide polymorphism
SSR: simple sequence repeat
SDP: strawberry diversity panel

